# Live-cell imaging of fetal microglia reveals time-dependent sex differences in their responsiveness to prenatal maternal stress

**DOI:** 10.64898/2026.10.23.761383

**Authors:** Alexandra Lawson, Matthew Rosin, Jessica M. Rosin

**Affiliations:** Neuroscience Graduate Program, Faculty of Medicine, University of British Columbia, Vancouver, British Columbia, V6T 1Z3, Canada; Department of Oral Biological and Medical Sciences, Faculty of Dentistry, University of British Columbia, Vancouver, British Columbia, V6T 1Z3, Canada; Life Sciences Institute, University of British Columbia, Vancouver, British Columbia, V6T 1Z3, Canada

## Abstract

Prenatal maternal stress is increasingly being linked to neurodevelopmental disorders (NDDs), and across diverse maternal stressors, immune dysregulation is emerging as a common feature. This suggests that fetal microglia may influence neurodevelopment by detecting changes in signaling from the intrauterine environment. Using a mouse model of prenatal maternal stress, where pregnant mice were exposed to cold from embryonic day 11.5 (E11.5) to E15.5, we previously demonstrated that fetal microglia show region-specific and morphology-dependent sex differences in their responsiveness to prenatal maternal stress using static analyses. Here, we extend these analyses by conducting live-cell imaging on the E15.5 hypothalamic paraventricular nucleus (PVN) to assess microglial morphology, motility, and phagocytic behavior in real-time under homeostatic conditions and in response to prenatal maternal cold stress exposure. Prenatal maternal cold stress induced sex- and time-specific changes in microglial morphology in the PVN, where female microglia showed increased branch complexity across the entire live-cell video recording, while male microglia only showed this response at early time-points. Interestingly, PVN microglia also showed baseline sex differences in phagocytic behavior, with male microglia showing higher phagocytic activity as compared to female microglia. In response to prenatal maternal cold stress, male microglia showed a decrease in completed phagocytic events in the PVN, whereas female microglia showed increased engagement in phagocytic activity with no change in completed phagocytic events. Microglial motility was also altered in the PVN of female embryos at early time-points in response to prenatal maternal cold stress, but no such changes were observed in males. Together, these data demonstrate that fetal microglia show sex differences in key physiologies both at baseline and in response to prenatal maternal cold stress. Through time-resolved assessment of microglial dynamics, our findings also suggest that microglial responsiveness to prenatal maternal cold stress differs between males and females, which may help to provide a cellular basis for sex differences observed across many NDDs.

## Background

Prenatal maternal stress is not a uniform exposure; different stressors engage distinct signaling cascades, producing different local signals within the fetal brain (1–12). Psychosocial stressors, for example, act predominantly through hypothalamic-pituitary-adrenal (HPA) axis activation and glucocorticoid signaling, whereas immune challenges more prominently engage maternal cytokine pathways (6,13–20). Among these stressors, cold stress is especially relevant to the hypothalamus, as exposure to cold stress during pregnancy induces coordinated molecular changes within the maternal hypothalamus, including altered corticosteroid-receptor signaling and inflammatory cascade activation, in addition to elevating maternal corticosterone levels and inducing placental inflammation (9,20,21).

The effects of prenatal maternal cold stress on fetal neurodevelopment have been examined in the embryonic day 15.5 (E15.5) hypothalamus, where single-cell RNA sequencing (scRNAseq) identified a population of fetal microglia located in direct contact with neural stem and progenitor cells (NSPCs) that were transcriptionally responsive to prenatal maternal cold stress (1). Exposure to prenatal maternal cold stress also drove a reduction in oxytocin (OXT) neurons in the paraventricular nucleus (PVN) and led to long-term alterations in social behaviors, both of which were reversed by microglial depletion, but only in male offspring (1). These findings establish that prenatal maternal cold stress produces sex-specific transcriptional changes in fetal hypothalamic microglia; however, it is important to acknowledge that transcriptional signatures only capture one dimension of microglial state. Fully characterizing microglial state requires more than one readout, as transcriptional, morphological, and functional approaches each resolve different aspects of microglial biology.

Transcriptional profiling across development has revealed that microglia follow diverse transcriptional trajectories, with distinct gene expression signatures and chromatin states characterizing successive developmental phases (1,22–29). Microglia are the most transcriptionally heterogeneous at early developmental time-points, and converge to a more consistent population later in life (22,25,29,30). For instance, developing microglia include subsets associated with high proliferative capacity, metabolic activity, and active engagement with developing neural elements (25). This makes understanding microglial dynamics at baseline and in response to prenatal maternal perturbations a moving target. Further, while transcriptional identity reveals differences in gene expression both at baseline and in response to stressors, it does not fully capture how microglia functionally behave within the tissue. Microglia are highly dynamic cells and are capable of migrating throughout the nervous system, signaling with neighboring cells, and engulfing cellular and synaptic material (1,3,24,31–40).

Morphology is one proxy for assessing microglial function in tissue (41,42). During phagocytosis, as well as other forms of cellular interaction, microglia rapidly reorganize their actin cytoskeleton to extend processes or form phagocytic cups (39,43–49). Because this same cytoskeleton remodeling underlies process extension, branching, and soma remodeling, morphological features such as branching complexity, process length, and soma size can reflect changes in microglial functional state (33,41,50–54). However, much like transcriptional analysis, morphology must be interpreted in a developmental, sex, and region-specific context.

In adults, homeostatic microglia are typically highly ramified, occupying distinct, non-overlapping territories (35,37,55,56). Under inflammatory or pathological conditions, mature microglia transition toward an ameboid morphology, marked by retraction of branches and condensation of the soma (54,57–60). Embryonic microglia, in contrast, display much broader morphological variability and a greater prevalence of ameboid forms under homeostatic conditions; this is thought to support migration throughout the brain and elevated baseline phagocytic activity (24,25,31,35,61,62). Consistent with transcriptomic data suggesting that microglia converge toward a more stable population over development, microglia progressively acquire a more mature, ramified phenotype throughout development (22,25,35,63).

Beyond developmental and context-specific variation, morphology is also not a definitive proxy for function. Functional changes can occur without obvious structural changes, and similar morphologies may reflect distinct underlying functional behaviors (41,42,64,65). Recent work combining spatial transcriptomics and immunofluorescent staining in young and aged mouse brains showed that microglial morphology captures biologically meaningful heterogeneity but does not correspond one-to-one with transcriptomically defined state, as a subpopulation of disease-associated-like microglia retained a highly ramified morphology (65). Thus, even in the adult brain, binary classifications of morphological state can fail to capture the complexity of microglial activities.

A major bottleneck to microglial analysis is scale, as accurately capturing microglial complexity often requires assessment across regions, conditions, and/or sexes (41,42,55,66–69). Two-dimensional (2D) based workflows can help address these issues using high-throughput and computationally efficient pipelines, enabling classification and morphologic quantification of large numbers of microglial cells relatively quickly (70–72). However, 2D representations compress process geometry and can miss morphological complexity that is only visible in three-dimensional (3D) approaches (70,73–75). Static approaches, however, whether 2D or 3D, share a fundamental limitation in that they capture microglial state at a single moment. Live-cell or four-dimensional (4D) imaging addresses this directly, enabling minute-scale quantification of branching dynamics, surveillance territory, and motility that static snapshots cannot (37,76–81). Live imaging in embryonic and early postnatal tissue has revealed dynamic microglial behaviors, including shifts in motility, surveillance territory, and process dynamics, that unfold on timescales of minutes and would be invisible to static approaches (5,50,76,79,82).

Despite the inherently dynamic nature of microglial behavior, live imaging of fetal microglia, particularly within the hypothalamic PVN and in the context of prenatal maternal stress, remains largely unexplored. Previously, we identified a morphological shift in male microglia from ameboid to ramified in the fetal PVN in response to prenatal maternal cold stress, in addition to morphology-dependent increases in both microglia-arginine vasopressin (AVP) neuron interactions and microglial phagocytosis, but only in males (83). Herein, we build on our static analyses by performing live-cell imaging on E15.5 microglia in the PVN of male and female embryos to study microglial dynamics in real-time, both at baseline and in response to prenatal maternal cold stress. Under homeostatic conditions, we uncover baseline sex differences in fetal microglial phagocytic behaviors, whereby male microglia show higher levels of engagement in phagocytic activity and overall significantly more completed phagocytic events in comparison to female microglia. Our live-cell imaging data also reveal sex- and time-dependent changes in fetal microglial morphology, motility, and phagocytosis in response to prenatal maternal cold stress, illustrating additional avenues through which microglia may contribute to sex differences in the onset, presentation, and/or prevalence of neurodevelopmental disorders (NDDs).

## Results

### Exposure to prenatal maternal cold stress does not alter microglial soma size

Prenatal maternal cold stress exposure has been shown to drive both sex-specific and microglia-dependent disruptions in the embryonic PVN (1). More recently, static analyses of the embryonic PVN has also demonstrated that microglia show morphology-dependent sex differences in their responsiveness to prenatal maternal stress (83). Although these findings highlight important sex differences in microglial responsiveness to maternal stress in the embryonic brain, they only assess microglial pathophysiology at a single moment in time. Microglial motility and process dynamics can shift without any accompanying change in morphology (41,42,65,84), and dynamic behaviors such as surveillance, migration, and real-time engulfment unfold on timescales of minutes to hours, an effect that fixed-tissue analysis cannot resolve (37,48,49,78,79,81,85) . Live-cell or 4D imaging of embryonic and early postnatal tissues has revealed dynamic behaviors including shifts in motility, surveillance territory, and process dynamics (5,37,48,76,79,82) . Therefore, given the inherently dynamic nature of microglia, conducting live imaging of microglia within the embryonic PVN, both at baseline and in response to prenatal maternal stress, will address the limitations of prior studies.

Herein, we utilized our previously established prenatal maternal cold stress model, whereby pregnant dams are subjected to 30 minutes of 4°C cold exposure daily from E11.5 to E15.5, to study the impact of prenatal maternal stress on microglial dynamics in the embryonic PVN. Live E15.5 *Cx3cr1^GFP/+^* hypothalamic vibratome sections containing the PVN were imaged across ∼6 hours (Figure S1A), with one frame captured every ∼8 min, such that each frame corresponded to one sampled time-point. Standardized exclusion criteria were applied to retain only viable, trackable cells, with dead cells (Figure S1B), cells with a bursting or non-contiguous soma (Figure S1C), and cells whose soma had adhered to the cut surface (Figure S1D), being excluded from downstream analysis. All remaining cells were quantified using *Imaris* software to assess microglial morphology, motility, and phagocytic activity over time (Figure S1E, E’).

We first assessed microglial soma volume in the E15.5 control and prenatal maternal cold stress male and female PVN. Interestingly, microglial cells exhibited considerable soma heterogeneity across all groups (Figure 1A). We quantified microglial soma volume at consistent intervals across the entire recording (approximately every hour) to obtain representative measures. When averaged across the full recording, microglial soma volume did not differ by sex or in response to prenatal maternal cold stress exposure (Figure 1B, Table S1; treatment, F_(1,126)_=1.591, p=0.2096; sex, F_(1,126)_=0.1451, p=0.7039; treatment×sex interaction, F_(1,126)_=1.111, p=0.2938). We next assessed soma volume at each of the individual sampled time-points, which were acquired at consistent intervals across the entire recording and allowed us to assess whether soma size changed across time. In males, prenatal maternal cold stress exposure did not alter microglial soma volume in the PVN at any time-point assessed (Figure 1C, Table S2; treatment, F_(1,65)_=2.183, p=0.1444; treatment×time interaction, F_(6,302)_=0.1148, p=0.9946); although microglial soma volume varied significantly across the recording period (Figure 1C, Table S2; time, F_(6,302)_=4.086, p=0.0006). Similarly in females, prenatal maternal cold stress exposure did not alter microglial soma volume in the PVN at any time-point analyzed (Figure 1D; Table S3; treatment, F_(1,61)_=0.0562, p=0.8134; treatment×time interaction, F_(6,272)_=1.463, p=0.1911); although, like males, microglial soma volume varied significantly over the recording period (Figure 1D; Table S3; time, F_(6,272)_=3.873, p=0.0010).

**Figure 1.**
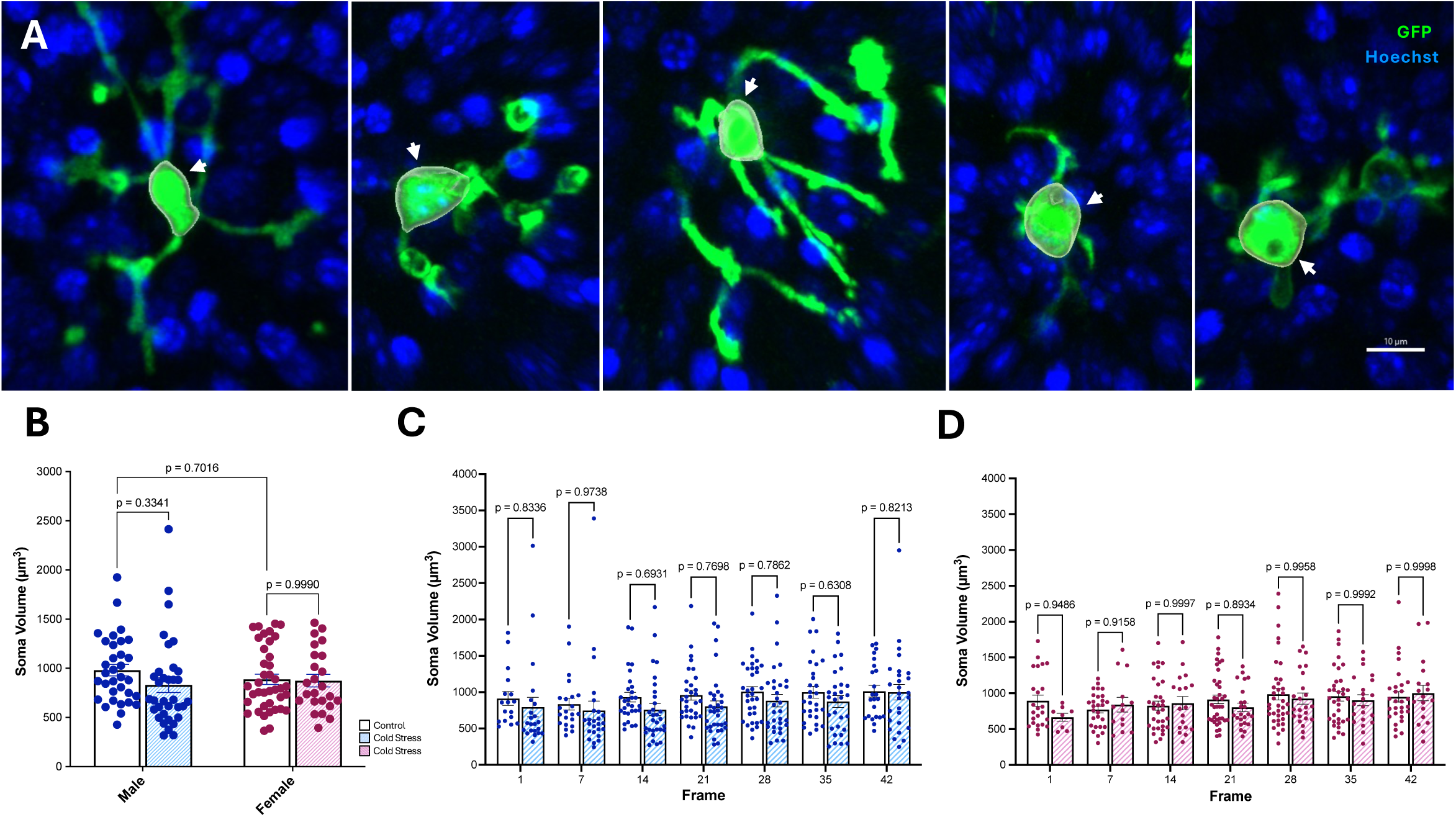
Microglial soma volume in the E15.5 control and cold stress PVN across time. (A) Representative images of microglial soma morphology in the E15.5 PVN. The soma is outlined in white and indicated by a white arrow. (B) Microglial soma volume averaged across time in the E15.5 control and cold stress male and female PVN (treatment, p=0.2096; sex, p=0.7039; treatment×sex interaction, p=0.2938). (C) Microglial soma volume by time-point in the E15.5 control and cold stress male PVN (treatment, p=0.1444; time, p=0.0006; treatment×time interaction, p=0.9946). (D) Microglial soma volume by time-point in the E15.5 control and cold stress female PVN (treatment, p=0.8134; time, p=0.0010; treatment×time interaction, p=0.1911). (B–D) Data are represented as means ± SEM and were analyzed by two-way ANOVA with Tukey-adjusted post hoc comparisons (B) or a mixed-effects ANOVA with Sidak-adjusted post hoc comparisons (C, D). Each dot represents an individual microglial cell (n=24-39) from N=3-5 independent embryos.

These findings indicate that microglial soma volume varies across time in the fetal PVN; however, microglial soma volume does not differ by sex or in response to prenatal maternal cold stress exposure over the course of the recording.

### Prenatal maternal cold stress drives sex- and time-dependent changes in microglial branching morphology in the fetal PVN

To study the impact of prenatal maternal cold stress on microglial morphology in real-time, we examined various aspects of microglial branching in the E15.5 PVN using *Imaris*, where several morphological features were extracted, including branch points, terminal points, Sholl intersections, and total branch length (Figure 2A). Using the same time-point sampling described in the microglial soma volume analysis, we initially noted that microglia appeared to generally exhibit greater branching complexity at early time-points relative to late time-points in both sexes (Figure 2B–I; Additional Files 1-4). While our live hypothalamic vibratome slices were rested for 1 hour prior to imaging to account for microglial reactivity in response to slicing the tissue, these results are consistent with evidence that microglia rapidly extend processes toward the cut surface in acute slice assays (86,87).

**Figure 2.**
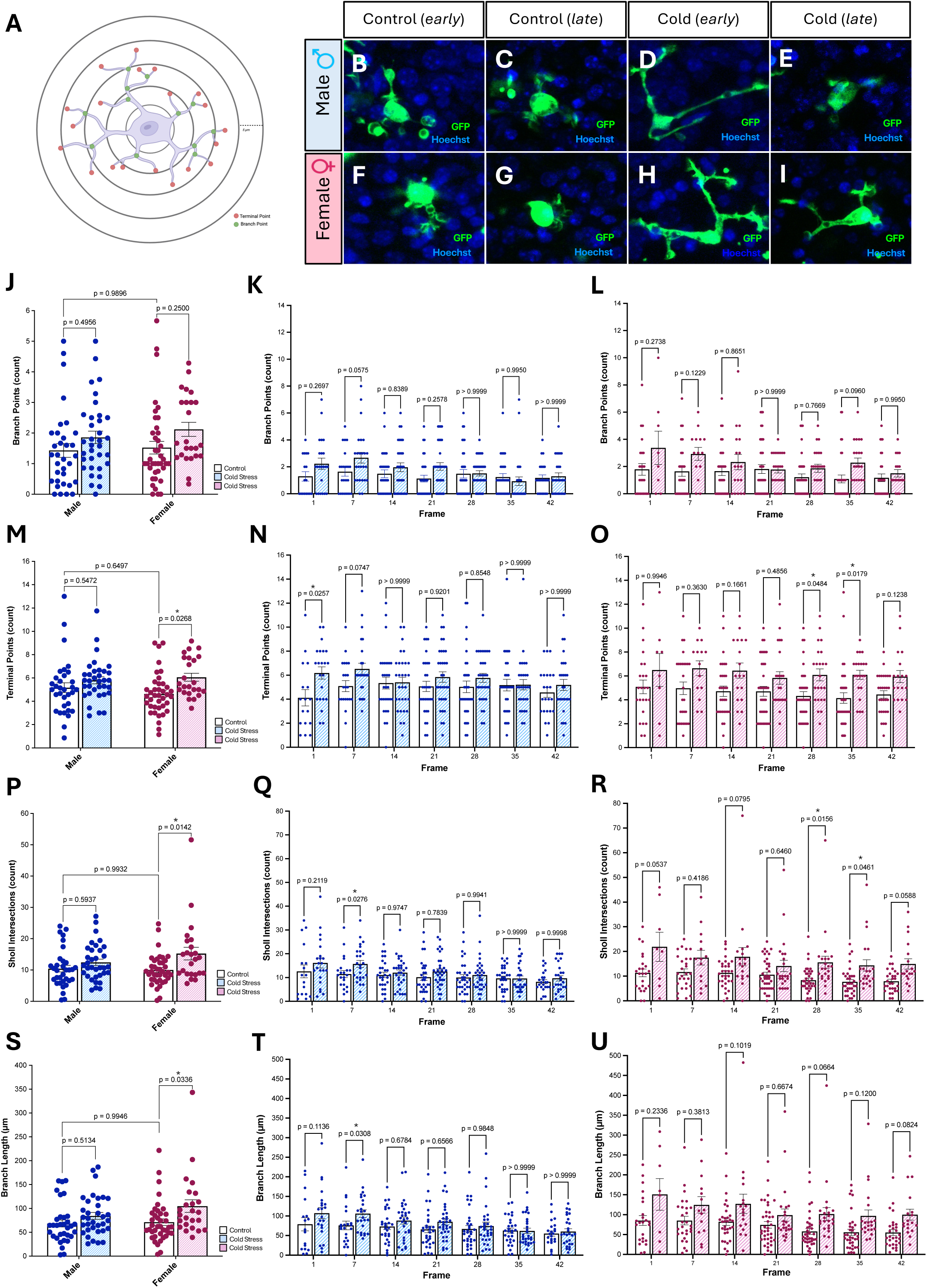
Microglial morphological branch complexity in the E15.5 control and cold stress PVN across time. (A) Microglial morphological measurements extracted using *Imaris*. (B–I) Representative images of microglia from control male (B, C), cold stress male (D, E), control female (F, G), and cold stress female (H, I) embryos at early and late time-points. (J) Microglial branch points across time in control and cold stress males and females (treatment, p=0.0219; sex, p=0.4252; treatment×sex interaction, p=0.6983). (K–L) Microglial branch points by time-point in control and cold stress males (K; treatment, p=0.1139; time, p=0.0008; treatment×time interaction, p=0.0249) and females (L; treatment, p=0.0271; time, p=0.0001; treatment×time interaction, p=0.1740). (M) Microglial terminal points across time in control and cold stress males and females (treatment, p=0.0035; sex, p=0.6965; treatment×sex interaction, p=0.2499). (N–O) Microglial terminal points by time-point in control and cold stress males (treatment, p=0.1302; time, p=0.3271; treatment×time interaction, p=0.0107) and females (O; treatment, p=0.0040; time, p=0.5502; treatment×time interaction, p=0.7445). (P) Microglial Sholl intersections across time in control and cold stress males and females (treatment, p=0.0025; sex, p=0.3140; treatment×sex interaction, p=0.1739). (Q–R) Microglial Sholl intersections by time-point in control and cold stress males (Q; treatment, p=0.1480; time, p<0.0001; treatment×time interaction, p=0.0193) and females (R; treatment, p=0.0040; time, p=0.0003; treatment×time interaction, p=0.4227) PVN. (S) Microglial total branch length across time in control and cold stress males and females (treatment, p=0.0038; sex, p=0.1676; treatment×sex interaction, p=0.2917). (T–U) Microglial total branch length by time-point in the control and cold stress males (T; treatment, p=0.1020; time, p<0.0001; treatment×time interaction, p=0.0182) and females (U; treatment, p=0.0099; time, p<0.0001; treatment×time interaction, p=0.7570). (J–U) Data are represented as means ± SEM and analyzed by two-way ANOVA with Tukey-adjusted post hoc (J, M, P, S) or a mixed-effects ANOVA with Sidak-adjusted post hoc (K–L, N–O, Q–R, T–U). Each dot represents an individual microglia cell (n=24-39) from N=3-5 independent embryos.

The number of microglial branch points, averaged across the full recording, was found to be significantly influenced by prenatal maternal cold stress exposure (Figure 2J, Table S4; treatment, F_(1,126)_=5.390, p=0.0219; sex, F_(1,126)_=0.6401, p=0.4252; treatment×sex interaction, F_(1,126)_=0.1509, p=0.6983). Although prenatal maternal cold stress appeared to broadly influence microglial branching complexity in the E15.5 PVN, this effect was not detectable within any individual comparisons as microglial branch points did not differ between control and prenatal maternal cold stress male (Figure 2J; p=0.4956) or female (Figure 2J; p=0.2500) embryos.

We next conducted time-specific analysis of microglial branch points in the E15.5 PVN, assessing differences in control and prenatal maternal cold stress embryos at each sampled time-point. In males, exposure to prenatal maternal cold stress drove a time-dependent change in microglial branch points across the recording period (Figure 2K, Table S5; treatment, F_(1,65)_=2.568, p=0.1139; treatment×time interaction, F_(6,302)_=2.453, p=0.0249). Although microglial branch points in prenatal maternal cold stress male embryos showed a trend towards an increase at frame 7 (Figure 2K; p=0.0575), a significant change in microglial branch points was not observed at any individual time-point between control and prenatal maternal cold stress males (Figure 2K). In females, exposure to prenatal maternal cold stress significantly altered microglial branch points across the recording period (Figure 2L, Table S6; treatment, F_(1,61)_=5.131, p=0.0271; treatment×time interaction, F_(6,272)_=1.513, p=0.1740). Overall effects suggested a persistent increase in microglial branch points in females in response to prenatal maternal cold stress, but this did not reach significance at any individual time-point assessed (Figure 2L). Interestingly, microglial branch points varied across time in both male (Figure 2K, Table S5; time, F_(6,302)_=3.940, p=0.0008) and female (Figure 2L, Table S6; time, F_(6,272)_=4.686, p=0.0001) embryos.

Next, we assessed microglial terminal points in the E15.5 control and prenatal maternal cold stress male and female PVN. Averaged across the full recording, prenatal maternal cold stress was found to significantly influence microglial terminal points (Figure 2M, Table S7; treatment, F_(1,126)_=8.859, p=0.0035; sex, F_(1,126)_=0.1529, p=0.6965; treatment×sex interaction, F_(1,126)_=1.336, p=0.2499). Further comparison demonstrated that while prenatal maternal cold stress did not alter microglial terminal points in the E15.5 male PVN (Figure 2M; p=0.5472), a significant increase in microglial terminal points was observed in females in response to prenatal maternal cold stress exposure (Figure 2M; p=0.0268). Interestingly, time-specific analysis of microglial terminal points in response to prenatal maternal cold stress revealed further sex differences. In males, exposure to prenatal maternal cold stress drove a time-dependent change in microglial terminal points across the recording period (Figure 2N, Table S8; treatment, F_(1,65)_=2.349, p=0.1302; treatment×time interaction, F_(6,302)_=2.831, p=0.0107). This effect was apparent at early time-points, with prenatal maternal cold stress males showing significantly more microglial terminal points than control males at frame 1 (Figure 2N; p=0.0257) and a trend towards an increase at frame 7 (Figure 2N; p=0.0747). In females, exposure to prenatal maternal cold stress drove a sustained, time-independent change in microglial terminal points across the recording period (Figure 2O, Table S9; treatment, F_(1,61)_=8.942, p=0.0040; treatment×time interaction, F_(6,272)_=0.5821, p=0.7445). This effect did not reach significance until later time-points, with prenatal maternal cold stress females showing significantly more microglial terminal points than control females at frame 28 (Figure 2O; p=0.0484) and frame 35 (Figure 2O; p=0.0179). In contrast to branch points, microglial terminal points did not vary by time across the recording period in either male (Figure 2N, Table S8; time, F_(6,302)_=1.610, p=0.3271) or female (Figure 2O, Table S9; time, F_(6,272)_=0.8266, p=0.5502) embryos.

As a measure of branching complexity, we assessed microglial Sholl intersections in the E15.5 control and prenatal maternal cold stress male and female PVN. When averaged across the full recording period, microglial Sholl intersections were found to be significantly impacted by prenatal maternal cold stress (Figure 2P, Table S10; treatment, F_(1,126)_=9.532, p=0.0025; sex, F_(1,126)_=1.022, p=0.3140; treatment×sex interaction, F_(1,126)_=1.870, p=0.1739). Further comparison demonstrated that while prenatal maternal cold stress did not alter microglial Sholl intersections in the E15.5 male PVN (Figure 2P; p=0.5937), a significant increase in microglial Sholl intersections was observed in females in response to prenatal maternal cold stress exposure (Figure 2P; p=0.0142). Intriguingly, time-specific analysis of Sholl intersections revealed time-dependent sex differences in microglial responses to prenatal maternal cold stress. In males, exposure to prenatal maternal cold stress drove a time-dependent change in microglial Sholl intersections across the recording period (Figure 2Q, Table S11; treatment, F_(1,65)_=2.144, p=0.1480; treatment×time interaction, F_(6,302)_=2.569, p=0.0193), which was most apparent at early time-points, with microglia in prenatal maternal cold stress males showing significantly more Sholl intersections than control males at frame 7 (Figure 2Q; p=0.0276). In females, prenatal maternal cold stress drove a sustained, time-independent increase in microglial Sholl intersections across the recording period (Figure 2R, Table S12; treatment, F_(1,61)_=8.971, p=0.0040; treatment×time interaction, F_(6,272)_=1.004, p=0.4227). This effect did not reach significance until later time-points, with prenatal maternal cold stress driving an increase in Sholl intersections at frame 28 (Figure 2R; p=0.0156) and frame 35 (Figure 2R; p=0.0461) in the female PVN. Interestingly, microglial Sholl intersections also varied across time in both male (Figure 2Q, Table S11; time, F_(6,302)_=9.671, p<0.0001) and female (Figure 2R, Table S12; time, F_(6,272)_=4.371, p=0.0003) embryos.

Finally, we assessed total microglial branch length in the E15.5 control and prenatal maternal cold stress male and female PVN. When averaged across the full recording period, total microglial branch length was found to be significantly impacted by prenatal maternal cold stress (Figure 2S, Table S13; treatment, F_(1,126)_=8.720, p=0.0038; sex, F_(1,126)_=1.927, p=0.1676; treatment×sex interaction, F_(1,126)_=1.121, p=0.2917). Further comparison demonstrated that while prenatal maternal cold stress did not alter total microglial branch length in the E15.5 male PVN (Figure 2S; p=0.5134), a significant increase in total microglial branch length was observed in females in response to prenatal maternal cold stress exposure (Figure 2S; p=0.0336). Moreover, time-specific analysis of total microglial branch length again revealed sex differences in response to prenatal maternal cold stress. In males, exposure to prenatal maternal cold stress drove a time-dependent change in total microglial branch length across the recording period (Figure 2T, Table S14; treatment, F_(1,65)_=2.751, p=0.1020; treatment×time interaction, F_(6,302)_=2.594, p=0.0182), with microglia from prenatal maternal cold stress males showing significantly greater total branch length than control males at frame 7 (Figure 2T; p=0.0308). In females, prenatal maternal cold stress drove a sustained increase in total microglial branch length across the recording period (Figure 2U, Table S15; treatment, F_(1,61)_=7.081, p=0.0099; treatment×time interaction, F_(6,272)_=0.5664, p=0.7570). Although microglial branch length in prenatal maternal cold stress female embryos showed a trend towards an increase at frame 28 (Figure 2U; p=0.0664), a significant change in microglial branch length was not observed at any individual time-point between control and prenatal maternal cold stress females (Figure 2U). Consistent with microglial branch points and microglial Sholl intersections, total microglial branch length varied across time in both male (Figure 2T, Table S14; time, F_(6,302)_=8.737, p<0.0001) and female (Figure 2U, Table S15; time, F_(6,272)_=7.071, p<0.0001) embryos.

Collectively, these morphological analyses reveal time-dependent sex differences in microglial branching morphology in response to prenatal maternal cold stress exposure in the fetal PVN. In male embryos, prenatal maternal cold stress was associated with a time-specific increase in branching complexity concentrated at early time-points that dissipated over the course of the recording. In contrast, in female embryos, prenatal maternal cold stress was associated with a sustained elevation in branching complexity that persisted across the recording period but did not reach significance until later time-points.

### Morphological clustering reveals differing time-specific microglial responses to prenatal maternal cold stress in the fetal PVN

To better understand dynamic changes in the proportion of microglia with distinct morphologies across time, and how this is impacted by prenatal maternal cold stress exposure, cluster-based analysis was performed on two representative time-points from the live-cell imaging data: frame 7 (∼1 hour into the recording) and frame 35 (∼4.6 hours into the recording), selected as early and late representative time-points, respectively, by adapting the morphology analysis toolset MicrogliaMorphology (70) for fetal microglial morphologic quantification (83). PCA revealed that the majority of variance was again captured by the first two PCs (Figure 3A, B). Given the limited number of microglia available per frame, cluster boundaries were less discrete than in our previous static analysis (83). Nevertheless, three morphologically distinct clusters emerged corresponding to similar ameboid, rod-like, and ramified subtypes as previously identified (Figure 3C, D) (83) . The ameboid cluster was characterized by high circularity and low branch number, the rod-like cluster by extended average branch length, low circularity, and sparse branch points, and the ramified cluster by abundant branch points and junctions alongside a large overall cell area (Figure 3D, E).

**Figure 3.**
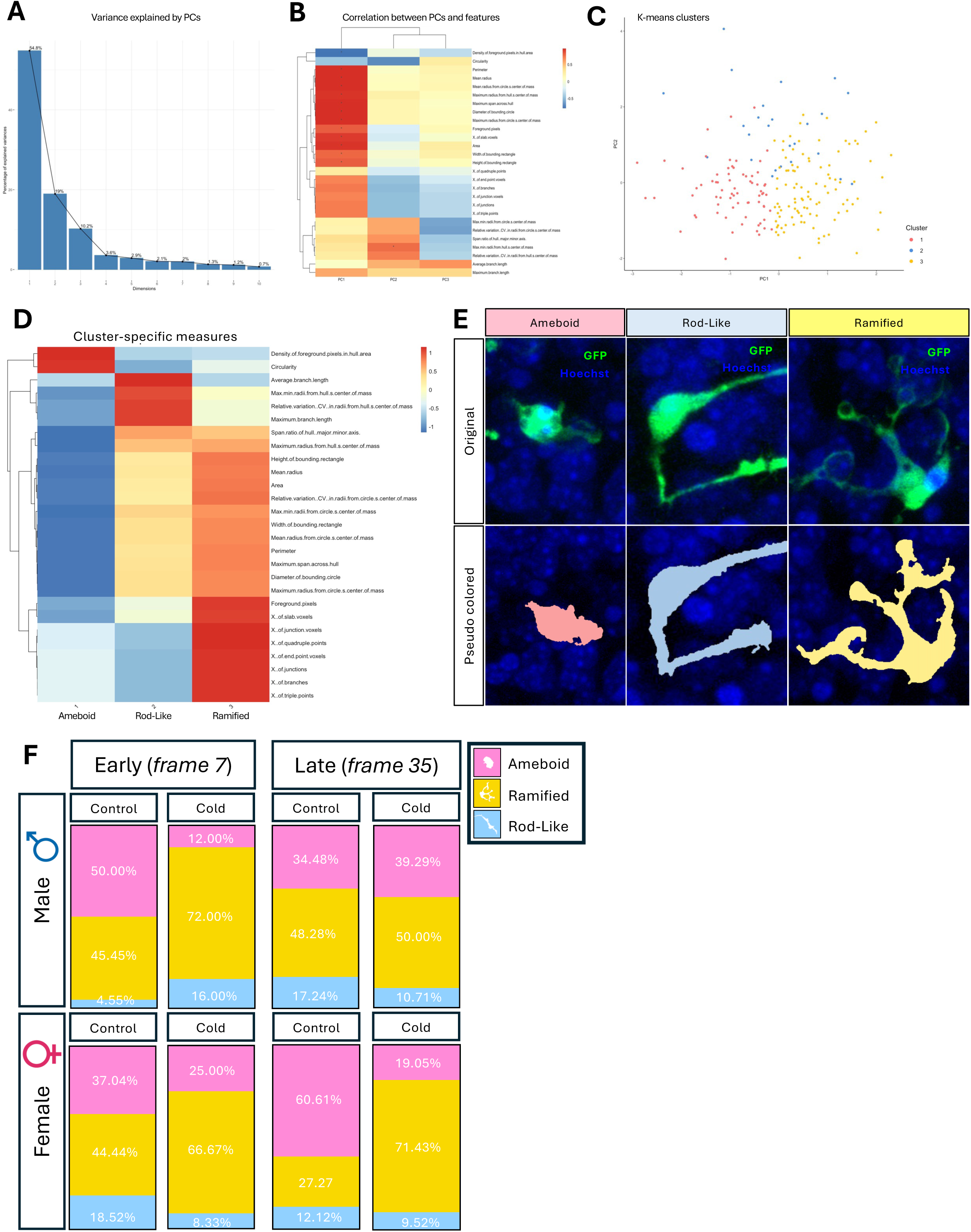
Unsupervised morphological clustering from live-cell imaging of microglia in the E15.5 PVN. (A) Elbow plot showing the proportion of variance explained by the first 10 principal components. (B) Heatmap of pairwise Pearson correlations among extracted microglial morphological features. (C) PCA of all microglial cells extracted. (D) Heatmap summarizing the relationship between individual morphological features and cluster identity. (E) Representative images of individual microglia from each identified cluster (ameboid, ramified, and rod-like) and their corresponding pseudo–colored reconstructions. (F) Proportion of each microglial cluster at early and late time-points in the E15.5 control and cold stress male and female PVN.

Due to low microglial counts per embryo at individual frames, in some cases as few as 2 to 3 microglia per embryo, proportional data were unsuitable for formal statistical comparison. Instead, microglia were pooled within each experimental group (control male, control female, cold stress male, and cold stress female) at each time-point (frame 7 and frame 35), and cluster proportions were examined descriptively. At the early time-point, a reduction in the proportion of ameboid microglia was observed in the E15.5 PVN in response to prenatal maternal cold stress in both male and female embryos (Figure 3F). In males, ameboid microglia comprised 50% of total microglia in control conditions, which dropped to 12% in prenatal maternal cold stress embryos (Figure 3F). A similar pattern was observed at the early time-point in females, where ameboid microglia occupied 37% in controls and dropped to 25% in prenatal maternal cold stress embryos (Figure 3F). At the late time-point, this pattern persisted in females but dissipated in males. In males, ameboid microglia occupied 34% of total microglia in controls compared to 39% in prenatal maternal cold stress embryos, illustrating almost no change in the proportion of ameboid microglia (Figure 3F). However, in females this pattern became even more pronounced compared with the early time-point, with controls exhibiting 60% ameboid microglia compared to 19% in prenatal maternal cold stress embryos (Figure 3F). In all cases, this shift from more to fewer ameboid microglia between control and prenatal maternal cold stress embryos both at early and late time-points was accompanied by an almost directly proportional increase in ramified microglia (Figure 3F).

This sex-divergent pattern is consistent with morphometric data observed from analysis of microglial branch complexity, where the effect of prenatal maternal cold stress on microglial morphology was sustained, if not amplified, across the recording period in females, but appeared to concentrate only at early time-points in males.

### Exposure to prenatal maternal cold stress drives sex-specific changes in microglia movement in the fetal PVN

Given that there is evidence that microglia respond to insults through movement (38,76,77), we next assessed microglial motility in the E15.5 control and prenatal maternal cold stress male and female PVN. Using the cell tracking feature in *Imaris*, we were able to document microglial movement across our entire recording (Figure 4A; Additional File 5). Total distance traveled by microglia in the E15.5 PVN differed broadly by sex, with no effect of prenatal maternal cold stress exposure (Figure 4B, Table S16; treatment, F_(1,127)_=1.927, p=0.1675; sex, F_(1,126)_=10.87, p=0.0013; treatment×sex interaction, F_(1,126)_=0.1136, p=0.7367). However, further examination of total distance traveled by microglia in the E15.5 PVN showed that it did not differ between control males and females (Figure 5B; p=0.1208).

**Figure 4.**
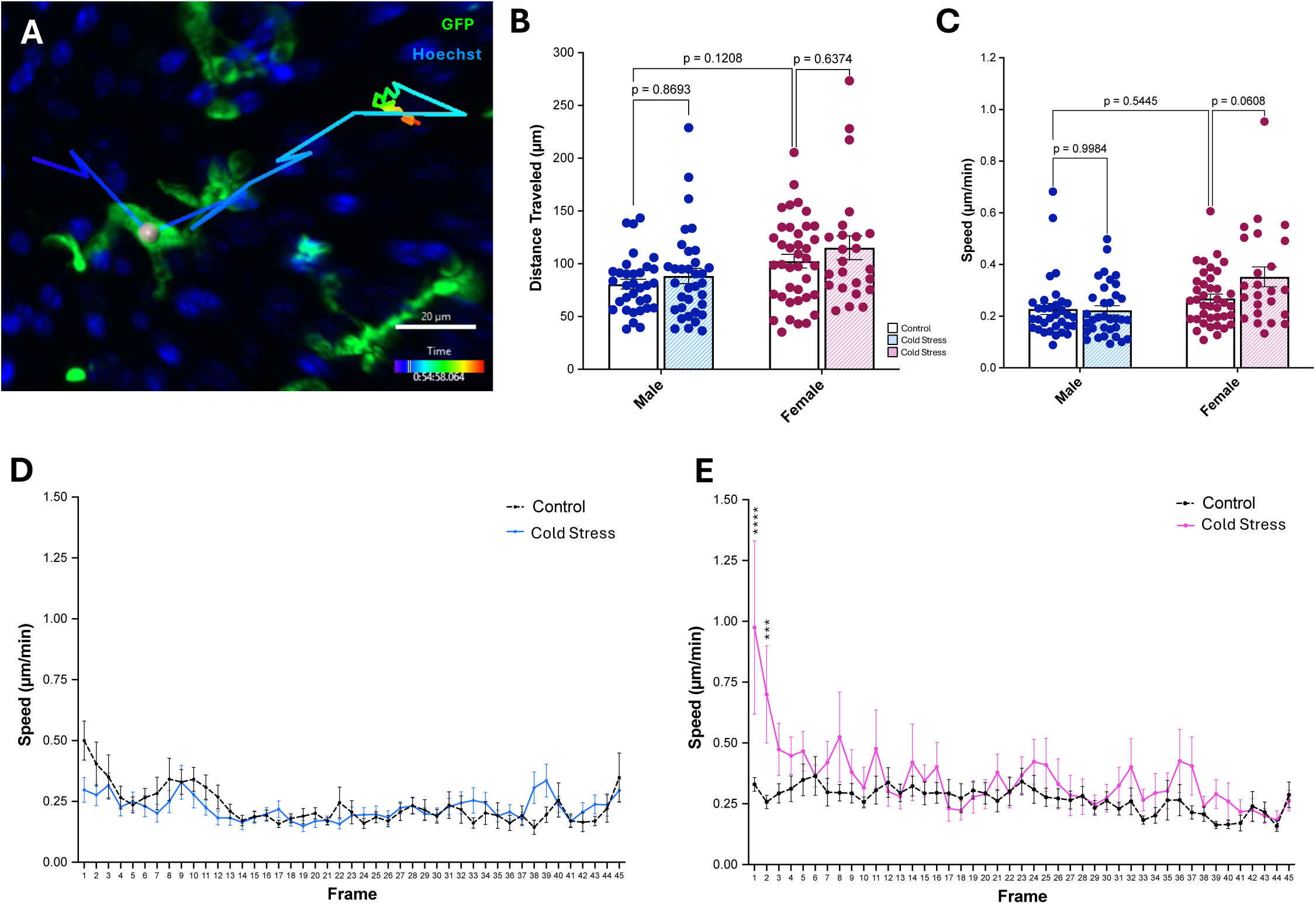
Microglial motility in the E15.5 control and cold stress PVN across time. (A) Representative image of *Imaris* tracking of microglial movement across an entire recording in the E15.5 PVN. (B) Total distance travelled by microglia in the E15.5 control and cold stress male and female PVN (treatment, p=0.1675; sex, p=0.0013; treatment×sex interaction, p=0.7367). (C) Microglial speed in the E15.5 control and cold stress male and female PVN (treatment, p=0.0867; sex, p=0.0003; treatment×sex interaction, p=0.0534). (D) Microglial speed by time-point in the E15.5 control and cold stress male PVN (treatment, p=0.5070; time, p<0.0001; treatment×time interaction, p=0.0023). (E) Microglial speed by time-point in the E15.5 control and cold stress female PVN (treatment, p=0.0080; time, p<0.0001; treatment×time interaction, p=0.0023). (B–E) Data are represented as means ± SEM and were analyzed by two-way ANOVA with Tukey-adjusted post hoc comparisons (B, C) or a mixed-effects ANOVA with Sidak-adjusted post hoc comparisons (D, E). (B, C) Each dot represents an individual microglial cell (n=24-39) from N=3-5 independent embryos.

**Figure 5.**
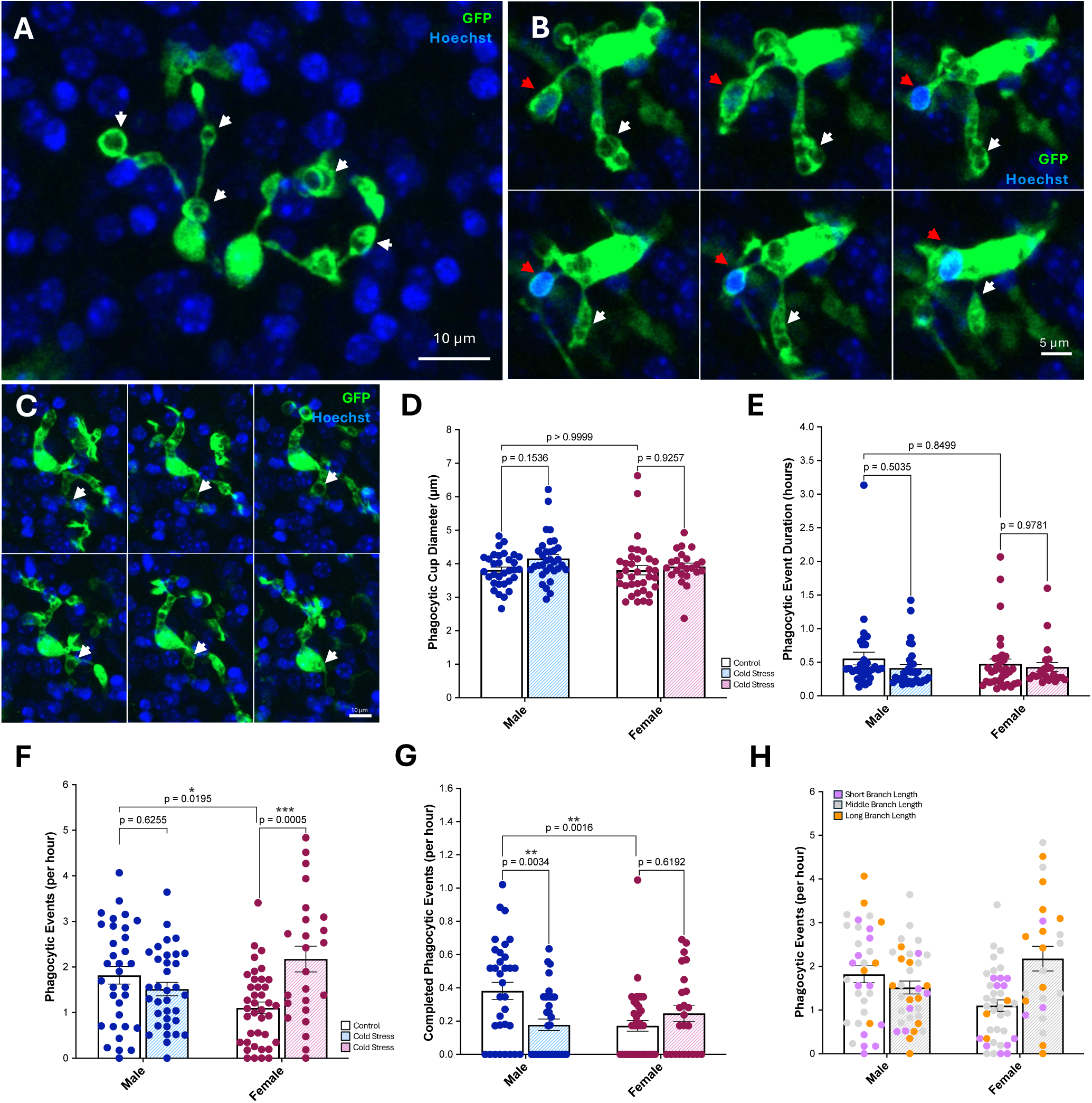
Microglial phagocytosis in the E15.5 control and cold stress PVN. (A) Representative image of microglia with multiple phagocytic cups in the E15.5 PVN. White arrows indicate phagocytic cups. (B–C) Representative images illustrating phagocytic events across 6 sequential frames in the E15.5 PVN. White arrows indicate an attempted event. Red arrows represent a completed event. (D) Microglial phagocytic cup diameter at maximum in the E15.5 control and cold stress male and female PVN (treatment, p=0.0609; sex, p=0.2997; treatment×sex interaction, p=0.3280). (E) Microglial phagocytic event duration in the E15.5 control and cold stress male and female PVN (treatment, p=0.2164; sex, p=0.6686; treatment×sex interaction, p=0.5143). (F) Total microglial phagocytic events per hour (attempted and completed) in the E15.5 control and cold stress male and female PVN (treatment, p=0.0369; sex, p=0.8748; treatment×sex interaction, p=0.0003). Phagocytic events were significantly higher in control males compared to control females (p=0.0195) and higher in cold stress females compared to control females (p=0.0005). (G) Completed microglial phagocytic events per hour in the E15.5 control and cold stress male and female PVN (treatment, p=0.1294; sex, p=0.0978; treatment×sex interaction, p=0.0013). Completed phagocytic events were significantly higher in control males compared to control females (p=0.0016) and in control males compared to cold stress males (p=0.0034). (H) Total microglial phagocytic events per hour (as displayed in F) and pseudo–coloured by morphology using total branch length quartiles: lowest 25% (purple), middle 50% (grey), and highest 25% (orange). (D–G) Data are represented as means ± SEM and were analyzed by two-way ANOVA with Tukey-adjusted post hoc comparisons. Each dot represents an individual microglial cell (n=24-39). Colour indicates sex (blue, male; pink, female), and shade within each colour denotes embryo (N=3-4).

As not all microglia were in the field of view for the entire recording, total distance traveled could be confounded by the duration a microglial cell was present rather than reflecting movement alone. Therefore, microglial speed was assessed in the E15.5 control and prenatal maternal cold stress male and female PVN. Microglial speed differed broadly by sex, with a trend toward an effect of prenatal maternal cold stress (Figure 4C, Table S17; treatment, F_(1,126)_=2.981, p=0.0867; sex, F_(1,126)_=13.82, p=0.0003; treatment×sex interaction, F_(1,126)_=3.804, p=0.0534). While microglial speed did not differ between control males and females (Figure 4C; p=0.5445) or following prenatal maternal cold stress in the E15.5 PVN of male embryos (Figure 4C; p=0.9984), a trend toward an increase in microglial speed was observed in prenatal maternal cold stress females relative to control females (Figure 4C; p=0.0608). Moreover, microglial speed was also found to be significantly higher in the E15.5 PVN of prenatal maternal cold stress females as compared to cold stress males (Table S17; p=0.0013).

Considering the time-specific differences in microglial morphology we observed, microglial speed was next assessed at each sampled time-point. As speed analysis required less manual processing than morphological reconstruction, all time-points were analyzed (i.e., every ∼8 min frame captured). In males, exposure to prenatal maternal cold stress drove a time-dependent change in microglial speed across the recording period (Figure 4D, Table S18; treatment, F_(1,65)_=0.4451, p=0.5070; treatment×time interaction, F_(44,2336)_=1.722, p=0.0023); although this did not resolve to any specific time-point (Figure 4D, Table S18). In females, exposure to prenatal maternal cold stress drove a time-dependent change in microglial speed across the recording period (Figure 4E, Table S19; treatment, F_(1,61)_=7.509, p=0.0080; treatment×time interaction, F_(44,2153)_=1.725, p=0.0023). This difference was concentrated at early time-points, with microglia from prenatal maternal cold stress females showing significantly higher speed than control females at frame 1 (Figure 4E; p<0.0001) and frame 2 (Figure 4E; p=0.0009). Interestingly, microglial speed varied across the recording in both male (Figure 4D, Table S18; time, F_(44,2336)_=4.046, p<0.0001) and female (Figure 4E, Table S19; time, F_(44,2153)_=3.524, p<0.0001) embryos.

Together, these findings suggest that exposure to prenatal maternal cold stress drives changes in microglial motility in the fetal PVN that differ between the sexes, with female microglia showing significantly increased speed at early time-points in response to prenatal maternal cold stress.

### Microglial phagocytic activity is sex-specific and shows divergent responses to prenatal maternal cold stress in the fetal PVN

Considering prior analyses illustrate that fetal microglia from male and female embryos display differing phagocytic responses to prenatal maternal cold stress, but were limited by static methodologies or the application of foreign bioparticles (83), we were interested in studying microglial engulfment behaviors in real-time in the E15.5 control and prenatal maternal cold stress male and female PVN using live-cell imaging. Microglial cells exhibited robust phagocytic activity, often presenting with multiple phagocytic cups simultaneously in the E15.5 PVN (Figure 5A). Considerable diversity was observed within this population of events: a subset of events, termed “complete”, were marked by both retraction into the soma and Hoechst+ nuclear staining (Figure 5B, red arrow; Additional File 6); other events had a phagocytic cup without retraction or Hoechst+ staining (Figure 5B, white arrow; Additional File 6); and others had a phagocytic cup and retracted into the soma but lacked Hoechst+ nuclear staining (Figure 5C; Additional File 7).

We first characterized the physical properties of all phagocytic cups in the E15.5 PVN, regardless of outcome. Microglial phagocytic cup diameter, at maximum, did not differ by sex or in response to prenatal maternal cold stress exposure (Figure 5D, Table S20; treatment, F_(1,119)_=3.579, p=0.0609; sex, F_(1,119)_=1.085, p=0.2997; treatment×sex interaction, F_(1,119)_=0.9646, p=0.3280). We next assessed phagocytic event duration in the E15.5 control and prenatal maternal cold stress male and female PVN. Microglial phagocytic event duration similarly did not differ by sex or in response to prenatal maternal cold stress exposure (Figure 5E, Table S21; treatment, F_(1,119)_=1.545, p=0.2164; sex, F_(1,119)_=0.1842, p=0.6686; treatment×sex interaction, F_(1,119)_=0.4276, p=0.5143).

Next, we assessed microglial phagocytic activity in the E15.5 control and prenatal maternal cold stress male and female PVN, again including all phagocytic events regardless of outcome. To account for the fact that not all microglia were present for the entire recording, we assessed this as the number of phagocytic events per hour. Interestingly, exposure to prenatal maternal cold stress had a significant influence on the frequency of microglial phagocytic events in a sex-dependent manner (Figure 5F, Table S22; treatment, F_(1,126)_=4.448, p=0.0369; sex, F_(1,126)_=0.02491, p=0.8748; treatment×sex interaction, F_(1,126)_=14.14, p=0.0003). Interestingly, microglia appeared to have significantly higher phagocytic activity in control males as compared to females (Figure 5F; p=0.0195). While microglial phagocytic activity did not differ in response to prenatal maternal cold stress in the E15.5 PVN of male embryos (Figure 5F; p=0.6255), exposure to prenatal maternal cold stress drove significantly higher phagocytic activity in female microglia in the E15.5 PVN (Figure 5F; p=0.0005).

As the preceding analysis captured any instance of phagocytic activity regardless of outcome, complete phagocytic events per hour were analyzed in the E15.5 control and prenatal maternal cold stress male and female PVN separately. Exposure to prenatal maternal cold stress had a significant influence on the frequency of complete microglial phagocytic events in a sex-dependent manner (Figure 5G, Table S23; treatment, F_(1,126)_=2.330, p=0.1294; sex, F_(1,126)_=2.781, p=0.0978; treatment×sex interaction, F_(1,126)_=10.85, p=0.0013). We again observed a baseline sex difference in microglial phagocytic activity, where microglia in control males had significantly more complete phagocytic events compared to control females (Figure 5G; p=0.0016). Intriguingly, exposure to prenatal maternal cold stress drove a significant reduction in completed phagocytic events by microglia in the E15.5 PVN of male embryos (Figure 5G; p=0.0034), while completed phagocytic events were comparable between control and prenatal maternal cold stress female microglia (Figure 5; p=0.6192).

Finally, to determine if we could associate a specific microglial morphology with a functional behavior, we examined whether total microglial branch length correlated with phagocytic event frequency in the E15.5 control and prenatal maternal cold stress male and female PVN. Microglial phagocytic events per hour were pseudo-colored by branch length, separated into quartiles: bottom 25% (purple), middle 50% (grey), and upper 25% (orange). However, no qualitative association between microglial branch length and phagocytic event frequency was observed in control or prenatal maternal cold stress male or female embryos (Figure 5H).

Together, these findings show that microglial phagocytic activity in the fetal PVN, both at baseline and in response to prenatal maternal cold stress, differs between the sexes. In contrast to prior static analyses, where correlations between specific microglial morphological phenotypes and phagocytic activity were observed in the fetal PVN (83), live-cell data suggests that 3D microglial branching morphology is not a reliable proxy for phagocytosis, and further that microglia across a range of morphologies are capable of phagocytic activity.

## Discussion

This study is the first to provide an in-depth characterization of fetal microglial morphology, motility, and phagocytic behaviors under homeostatic conditions and in response to prenatal maternal cold stress exposure using live-cell imaging. Through careful examination of the E15.5 hypothalamic PVN, we demonstrate that fetal microglia display sex- and time-dependent differences in microglial morphology in response to prenatal maternal cold stress. Microglia from male embryos showed early increases in branching complexity, whereas microglia from female embryos showed a more sustained increase in branch complexity across the entire live-cell imaging recordings, but this often did not reach significance until later time-points. In response to prenatal maternal cold stress exposure, time-dependent sex differences in microglial motility were also observed in the E15.5 PVN, where only female microglia showed increased motility at early time-points. Despite a lack of sex differences in microglial morphology at baseline, analysis of the E15.5 PVN revealed basal sex differences in microglial phagocytic behavior, where male microglia displayed both increased engagement in phagocytic activity and elevated completed phagocytic events compared to female microglia. Intriguingly, prenatal maternal cold stress also drove distinct microglial phagocytic responses that differed between the sexes, where female microglia showed heightened engagement in phagocytic activity and male microglia showed a decrease in completed phagocytic events. Together, these findings indicate that prenatal maternal cold stress does not produce a uniform microglial response but instead causes sex- and time-dependent changes in fetal microglia morphology, motility, and phagocytic behaviors, highlighting the importance of studying microglial dynamics both in real-time and across time.

Microglia are not static even under homeostatic conditions. Real-time imaging has shown that microglia continuously survey their environment by extending and retracting their processes (37,48,76,88,89), and rapidly adjust this behavior in response to perturbation of their surroundings (5,37,48,76–78). For example, *in vivo* two-photon imaging of microglia in the somatosensory cortex of adult mice revealed changes in surveillance and morphology during sleep-wake states and during sleep deprivation (78). Because such dynamics can unfold over minutes to hours, they are often missed in single time-point or static approaches. To capture this dimension, we utilized live-cell imaging to visualize microglial dynamics in the E15.5 PVN over a 6-hour period. Interestingly, and aligned with prior findings (83), we observed no sex differences in microglial morphology under homeostatic conditions. However, in response to prenatal maternal cold stress, and across multiple morphological measures (i.e., terminal points, Sholl intersections, and total branch length), we observed sex- and time-dependent changes in microglial morphology. Indeed, in response to prenatal maternal cold stress, microglia in the E15.5 male PVN exhibited an increase in branching complexity concentrated at early time-points, which resolved over the course of the 6-hour recording period. This contrasts with microglia in the E15.5 female PVN, which exhibited a sustained increase in microglial branching complexity when averaged across the 6-hour experiment, with statistically significant increases in branching complexity concentrated in the latter half of the recording. Depending on the time-point sampled, had the tissue been examined at only a single time-point, the male and female responses to prenatal maternal cold stress might have appeared similar, sex-specific or might have been missed entirely. This interpretation is supported by our own microglial morphology clustering analysis of an early and late frame from the 6-hour live-cell imaging recording. By applying an adapted version of the morphology analysis toolset MicrogliaMorphology (70) for fetal microglial morphologic analysis (83) at an early (frame 7) and late (frame 35) time-point, we observe a shift in male microglia towards a more ramified morphology at the early but not late time-point in response to prenatal maternal cold stress, whereas in females this shift was present at both early and late time-points. This raises the possibility that previous static analysis of microglial morphology in the E15.5 PVN (83) may have only captured microglia at a stage comparable to the early time-points of the live-cell recording, and underscores the importance of considering microglial dynamics and trajectories across time in the fetal brain. Further, while embryonic microglial dynamics have been studied through live-cell imaging in the embryonic mouse cortex (31,90) and hypothalamus (50,82), and extensively in the zebrafish embryo (48,49,79), none of this work was stratified by sex. Our finding that male and female microglia differ in the trajectories of their response to prenatal maternal cold stress in the fetal PVN is consistent with broader evidence for sex differences in microglial function and behavior across development (1,10,16,33,35,55,64,83,91,92), and suggests that the timing of these differences may matter as much as their presence or magnitude.

Importantly, these sex- and time-dependent differences in microglial morphology could be a result of several non-exclusive factors, including tissue preparation, the lasting programming effects of prenatal maternal cold stress, and the metabolic demands placed on microglia. To begin, microglia in *ex vivo* brain slices have been shown to migrate toward the cut surface and undergo time-dependent phenotypic changes in response to this insult (86,93). Although we incubated our sliced tissue for one-hour prior to the start of imaging to mitigate some of these acute responses, microglia were still observed interacting with the cut surface throughout the recording, and this insult likely shaped baseline dynamics. However, because all sexes and treatment groups experienced the same preparation, divergent microglial trajectories are unlikely to reflect the slice procedure alone. Instead, the fact that microglia from male and female embryos followed distinct time courses in their response, diverging as the recording progressed, suggests that sex and prenatal maternal cold stress shaped how microglia responded across time.

One explanation for these divergent trajectories in microglial response is that prenatal maternal cold stress may alter how microglia respond to subsequent challenges—a phenomenon often termed priming, where a prior perturbation durably reshapes microglial state and can produce either a heightened or a blunted response to later stressors or insults (94,95). The direction and magnitude of this response are not fixed, varying with the nature and duration of the initial insult and the brain region examined (96). For example, following maternal immune activation (MIA), microglia from offspring whose mothers experienced MIA had persistent decreases (blunting) in immune reactivity throughout development and replacing these perturbed microglial cells with naïve cells rescued these downstream phenotypes (97). In contrast, in adult female mice, prenatal restraint stress heightened the microglial and inflammatory response of the hippocampus to a later immune challenge (98). Adding to this complexity, responses are not always unidirectional. For instance, in an adult chronic-stress model, prior stress reshaped how microglia responded to a later acute stressor, producing either a heightened or a blunted response depending on the duration and intensity of the initial stressor (99). In our study, prenatal maternal cold stress may similarly have left microglia from male and female embryos in different states of responsivity, shaping how each responded over time during the imaging period.

Although priming can act through many molecular and transcriptional changes (100), one candidate mechanism for the sex-divergent trajectories we observed is metabolic demand. Microglial surveillance and phagocytosis are energetically costly, and microglial reactivity and changes in branch dynamics require metabolic reprogramming to increase ATP production (101,102). Interestingly, recent *in vivo* imaging has shown that microglial processes performing minute-to-minute surveillance and acute responses largely lack mitochondria, with mitochondrial support recruited only later (103). This implies that rapid, early process dynamics and sustained responses draw on distinct metabolic sources, with prolonged activity placing greater demand on mitochondrial metabolism. Our static findings of a shift from ameboid to ramified morphology within the male PVN, together with evidence that hypothalamic microglia from male embryos mount a greater transcriptional response to prenatal maternal cold stress as compared to female embryos (1), suggest that prenatal maternal stress heightens microglial reactivity in male embryos, potentially to a greater degree than in females. If already operating near their metabolic limits, microglia from prenatal maternal cold stress male embryos may mount an early response to the acute slice challenge but lack the metabolic reserves to sustain this response. In contrast, microglia from prenatal maternal cold stress female embryos may also be primed yet retain sufficient metabolic reserves to support a sustained response. In this framework, exposure to prenatal maternal stress may produce sex differences in microglial reactivity and consequently in the metabolic reserves available to meet a subsequent challenge.

Alternatively, reduced microglial process complexity could reflect declining cell health during imaging rather than a change in microglial state. Dying microglia retract their processes and round up before death (104), a morphology that overlaps with that of a healthy ameboid microglia. Accordingly, branching complexity alone cannot distinguish a dying microglial cell from a reactive one. This is relevant because sustained activation drives mitochondrial dysfunction and reactive oxygen species (ROS) production, increasing susceptibility to apoptosis (105,106), while live imaging itself generates additional ROS that can impair cell physiology and cause death (107,108). Microglia from male embryos, if already generating ROS through a sustained, cold-stress-driven response, may therefore have been less able to tolerate the added burden of acute slice injury. The accelerated decline in branching complexity we observed in microglia from prenatal maternal cold stress male embryos could thus reflect reduced viability, a genuine change in state or both. While assessing fetal microglial dynamics using intravital imaging of externally immobilized embryos or para-uterine imaging of a stabilized embryo within the abdomen of the pregnant mouse (109–111) may yield differing results, as they do not involve a slice insult, microglia are highly sensitive to the drugs and anesthetics used to image embryos in a living dam (112–114) and these technologies are often best suited to structures like the cortex, not structures deep in the brain such as the hypothalamic PVN. Therefore, there are different pros and cons for using each individual experimental methodology and all of them need to be considered when analyzing the datasets resulting from live-cell imaging.

Beyond microglial morphology, we also examined microglial movement. Unlike adult microglia, which are largely stationary and survey their environment by extending and retracting their processes, fetal microglia are migratory, moving through the brain as they colonize various regions (24,25,31,35,61,62). Microglia also rapidly migrate toward sites of injury or tissue damage (77,87); therefore, movement provides an additional functional readout distinct from morphology and phagocytosis. When averaged across the 6-hour recording, we observed no significant differences in microglial movement in the E15.5 control and prenatal maternal cold stress male and female PVN. However, at the earliest time-points of the recording (frames 1 and 2) female microglia showed increased movement in response to prenatal maternal cold stress. This early increase in movement may reflect an enhanced or accelerated migratory response to the acute slice insult, as early-life stress is known to modulate microglial behavior. For instance, *in vivo* two-photon imaging in the adult mouse somatosensory cortex has shown increased motility of microglial processes following early-life maternal separation (115). Importantly, this early, female-specific increase in movement observed in our study may represent a sex-specific component of the microglial response, possibly being shaped by both the lasting priming effects of prenatal maternal cold stress exposure and the acute slice insult.

Our live-cell imaging analysis of phagocytosis added further complexity to these sex- and time-dependent microglial responses. Phagocytosis is a dynamic, multi-step process, progressing from the detection of a target, through the extension of processes, formation of a phagocytic cup, engulfment, and eventual digestion of the engulfed material (116). Live-cell imaging allows us to resolve not only a single snapshot but also the dynamics of this complex process (37,76–81). We first quantified the diameter and duration of microglial phagocytic events and observed no differences across sexes or in response to prenatal maternal cold stress. In principle, the size and duration of a phagocytic event carry information about what is being engulfed, as the size and shape of a target are likely to influence both the dimensions of the phagocytic cup and the duration of engulfment (117). The absence of differences therefore suggests that prenatal maternal cold stress did not alter the physical characteristics of these events, or by extension, the broad nature of the cells or material being engulfed.

Size and duration, however, capture only one part of a microglial phagocytic event. Previous work has additionally distinguished microglial phagocytic events by whether they are carried through to completion. In the developing zebrafish, for instance, microglial clearance of apoptotic neurons, termed “efferocytosis”, has been shown to involve the extension of multiple branches and phagocytic cups, of which typically only one results in a successful engulfment event at a time, with attempts sometimes aborted before completion (118). A comparable distinction has been noted in recent imaging of microglial responses to dying neurons: in the mouse cortex, microglial “engagement”—the ability of a microglial cell to find, move toward, and contact a dying neuron, including formation of an initial phagocytic cup, and microglial “clearance”, the successful phagocytosis and complete removal of the apoptotic cell, have been separately assessed (119). Other similar work has described microglial “trogocytosis”, the partial nibbling of targets rather than whole-cell or whole-synapse engulfment (39). With this in mind, we distinguished two categories of microglial phagocytic events. A microglial phagocytic event was counted whenever a microglial cell formed a phagocytic cup, regardless of whether it contained Hoechst+ nuclear material or was retracted into the microglial soma. In contrast, a “completed” microglial phagocytic event was counted only when a phagocytic cup was formed, contained Hoechst+ nuclear material, and was visibly drawn into the microglial soma. Total microglial phagocytic events therefore captured any engulfment behaviour, whereas completed events captured a narrower subset of these phagocytic activities.

When these phagocytic categories were applied to microglia analyzed from our live-cell imaging of the E15.5 PVN, we observed sex differences, both at baseline and in response to prenatal maternal cold stress. Under homeostatic conditions, control male microglia showed an increased rate of both total and completed microglial phagocytic events compared to control female microglia. This is notable, as it shows that even in the absence of sex differences in microglial morphology at baseline, functional sex differences still exist, adding to evidence that microglial morphology and function do not always correlate in a one-to-one manner (41,84). Additionally, these baseline differences in microglial phagocytic activity may be relevant to sex-specific effects of stress: if microglia from male and female embryos have divergent microglia activities under homeostatic conditions, they may also respond differently to challenge. Indeed, we observed distinct microglial responses to prenatal maternal cold stress in each sex, as well as differences across our total and complete microglial phagocytosis measures. For instance, exposure to prenatal maternal cold stress drove increased rates of total microglial phagocytic events in female embryos but did not drive any changes in male embryos. In contrast, when stratifying by completed microglial phagocytic events, we saw no change in female embryos in response to prenatal maternal cold stress, but a significant reduction in completed microglial phagocytic events in prenatal maternal cold stress males relative to control males. This suggests that microglia may engage with phagocytic targets differently depending on sex. Because microglia engulf a wide range of substrates, including apoptotic cells, viable neural progenitors, synapses, axons, and myelin (31,116,120,121), and have been shown to engage in partial or incomplete phagocytic events (39,118,119), treating all microglial phagocytic events as equal may obscure finer distinctions in target and downstream effects. Therefore, our findings illustrate both baseline sex differences in microglial phagocytic activity and sex-specific microglial phagocytic responses to prenatal maternal cold stress, highlighting the importance of considering sex as well as the finer intricacies associated with microglial phagocytic behavior.

Intriguingly, we also observed many cases of unique microglial behaviors, such as microglia-microglia interactions (Figure S2; Additional File 8). Observations like this reveal a plethora of microglial behaviors that can be assessed using live-cell imaging, the complexity in the activities of fetal microglia, and the potential importance of considering these types of unique and dynamic movements when studying the developing brain.

## Conclusions

This study provides some of the first characterizations of microglial dynamics in the E15.5 hypothalamus using live-cell imaging, resolving microglial morphology, motility, and phagocytic behaviors in both male and female embryos. By capturing these key microglial physiologies as they unfold over time, we show the diverse trajectories of fetal microglial responses to prenatal maternal stress. Additionally, divergent responses between total and completed microglial phagocytic events, both at baseline and following prenatal maternal stress exposure, illustrate that the types of phagocytic activities, not just total levels of phagocytosis, are worth investigating further. More broadly, these results reveal how heterogeneous fetal microglial responses to prenatal maternal stress can be, varying across sex and time, and involving changes in functional behavior that are not directly mirrored morphologically. Accounting for and further investigating this heterogeneity will be central to understanding how fetal microglia translate a perturbed intrauterine environment into lasting changes in offspring neurodevelopment and behavior.

## Methods

Datasets and additional *Imaris* protocols can be found on the GitHub: https://github.com/RosinLabUBC/Live-Cell-Microglia-Cold-Stress-Project-AL.git

### Experimental model

All experimental work conducted falls under the rules and restrictions outlined in Biosafety Protocol B21-0133, which received approval from the University of British Columbia. Animal work was carried out in accordance with the guidelines and regulations of the Canadian Council on Animal Care and received prior approval from the University of British Columbia’s Animal Care Committee (protocols A21-0170 and A21-0171).

To generate heterozygous *Cx3cr1^eGFP^* embryos for live-cell experiments, homozygous *Cx3cr1^eGFP/eGFP^* mice (*B6.129P2(Cg)-Cx3cr1^tm1Litt^/J*; The Jackson Laboratory 005582), which express green fluorescent protein (eGFP) under control of the endogenous *Cx3cr1* locus, were crossed with C57BL/6J mice. Breeding pairs consisted of either homozygous *Cx3cr1^eGFP/eGFP^* males with wild-type (WT) females or WT males with homozygous *Cx3cr1^eGFP/eGFP^* females, resulting in exclusively heterozygous *Cx3cr1^eGFP/+^*embryos. All experiments were blinded to sex.

### Mouse handling and maternal cold stress paradigm

In the morning, following the setup of a breeding pair, vaginal plugs were checked. Embryonic day 0.5 (E0.5) was designated as the day of initial plug detection. During pregnancy, pregnant mice received standard chow (LabDiet PicoLab Rodent Diet 20) and water *ad libitum* and were housed individually. Pregnant mice were assigned to control or cold stress conditions randomly. From E11.5 to E15.5, pregnant mice in the cold stress group were placed for 30 minutes daily in a 4°C walk-in refrigerator between 8:30am and 9:00am. During this time pregnant mice remained in their home cages with *ad libitum* access to standard chow (LabDiet PicoLab Rodent Diet 20) and water. On E15.5, to facilitate embryo collection, pregnant mice were anaesthetized with isoflurane, immediately euthanized by cervical dislocation, and decapitated. All mouse embryos were genotyped for sex.

### DNA extraction and genotyping

Embryonic sex was determined by PCR genotyping as previously described in Lawson et al. (2026) (83). In brief, DNA was extracted from E15.5 embryonic tail tissue by incubating overnight at 56°C in an extraction buffer supplemented with proteinase K (New England Biolabs P8107S). Genomic DNA was then isolated from digested tissue by saturated NaCl and isopropanol precipitation. DNA pellets were washed with 70% ethanol, dried at 37°C, resuspended in Tris-EDTA buffer, and incubated at 68°C for 1 hr. Polymerase chain reaction (PCR) was performed by combining genomic DNA with OneTaq Quick-Load 2X Master Mix (New England Biolabs M0486S) and 1 µM primers targeting the sex-linked genes *Sly* and *Xlr* (122) on an Applied Biosystems™ MiniAmp™ Thermal Cycler (Applied Biosystems), using an initial denaturation at 95°C for 3 min followed by 35 cycles of denaturation at 95°C for 30 sec, annealing at 57°C for 30 sec, and extension at 72°C for 30 sec, then through a final extension at 72°C for 5 min. Amplified PCR products were run on a 1.5% agarose gel in TAE buffer, visualized with SmartGlow pre-stain, and sized against a GeneRuler Ready-to-Use 100 bp DNA Ladder (Thermo Fisher Scientific SM0243).

### Live-cell imaging of microglia

Live-cell methods were adapted from Rosin et al. (81), and procedural details are provided in the methodology chapter by Lawson et al. (123). In brief, immediately following cold stress exposure, pregnant dams were euthanized and E15.5 embryos were placed in ice-cold PBS (Gibco 10010-023, Thermo Fisher Scientific). E15.5 brains were dissected in ice-cold PBS and embedded in UltraPure LMP 4% agarose (Invitrogen 16520050, Thermo Fisher Scientific) diluted in PBS. Coronal sections were cut at 350 µm thickness using a Leica VT1200S vibratome at a cutting speed of 0.3 mm/s and amplitude of 1.00 mm. Hypothalamic PVN sections (section 7) were identified based on landmarks and the characteristic aggregation of microglia within this region, and placed on top of 0.4 μm, 30 mm Millicell cell culture inserts (Millipore PICM0RG50) within 40 mm glass-bottom Petri dishes (Ted Pella Inc. 14026-20) containing 1.5 mL of warmed live-cell media with Hoechst (1:10000, Invitrogen H3570) and incubated in a humidified chamber for 1 hr (5% CO_2_, 37°C, humidified) to allow penetration throughout the slice. Live-cell media was prepared fresh prior to each experiment to a total volume of 8 mL, consisting of 4.6 mL DMEM (Gibco 11965-092, Thermo Fisher Scientific), 2.3 mL F12 (Gibco 11765-054, Thermo Fisher Scientific), 0.4 mL FBS, 0.4 mL horse serum (Gibco 26050-070, Thermo Fisher Scientific), 0.16 mL B27 supplement (Gibco 17504-044, Thermo Fisher Scientific), 0.08 mL N2 supplement (Gibco 17502-048, Thermo Fisher Scientific), and 0.08 mL GlutaMAX (Gibco 35050-061, Thermo Fisher Scientific). Following incubation, live-cell media containing Hoechst was removed and 1.5 mL of fresh warm live-cell media without Hoechst was added to the culture dish.

Live-cell imaging was performed using a Zeiss LSM 900 confocal microscope equipped with an environmental control chamber and 40×/0.6 NA long working distance air objective. The chamber was equilibrated to 37°C with 5% CO and 90% humidity. The heated lid was set to 40°C. Z-stacks were acquired from the basal surface of the slice (i.e., the surface in contact with the membrane and closest to the objective) to a depth of approximately 50 µm. Imaging was performed using a pinhole size of 1 AU (69 µm), a digital zoom of 0.5x, and an optimal Z-step size of ∼1 µm. Images were acquired with a voxel size of 0.137 µm × 0.137 µm × 1.03 µm (X×Y×Z). To increase the number of microglia captured per recording, the Zeiss *Tiles* function was used to acquire a two-tile region with 10% overlap encompassing the PVN. Each tile consisted of 2328 × 2328 pixels collected at 16-bit depth with no signal averaging. Acquisition parameters included an LSM scan speed of 8 (maximum), a pixel dwell time of 0.33 µsec, a frame time of 2.12 sec, bidirectional scanning, and a detector gain of 650 V. Laser power was optimized independently for Hoechst (3.5%) and GFP (4.0%) and configured as separate acquisition tracks using 405 nm and 488 nm solid-state lasers, respectively. Fluorescence emission was detected using gallium arsenide phosphide photomultiplier tube (GaAsP-PMT) detectors, collecting the full emission spectrum for each channel.

Time-lapse imaging was performed by acquiring 45 consecutive frames of the complete X×Y×Z two-tile volume at approximately 8-min intervals, corresponding to the time required to complete acquisition of a single two-tile Z-stack. This resulted in a total imaging duration of approximately 6 hrs. Each time series was initiated immediately upon completion of the previous acquisition, with no downtime, to capture microglial dynamics occurring on the scale of minutes. Once completed, videos were further processed using the ZEN 3.8 software to perform *LSM Plus Processing* (deconvolution) and stitch the two tiles together.

### Live-cell analysis of microglia using *Imaris*

Procedural details can be found in Lawson et al. (123). Using *Imaris* (*Imaris* v10.2, Oxford Instruments)*, .czi* files were converted to *.ims* using the *Imaris File Converter* plugin and analyzed to identify viable microglia. A cell was considered “viable” if (i) its soma was entirely within the field of view for ≥50% of captured frames and (ii) its soma remained intact (i.e., not ruptured, not visibly adhered/flattened against the surface, and not truncated by the image borders) for ≥50% of captured frames. Dead cells were excluded and were distinguished by a round, enlarged soma lacking processes, 3D shape, with absent branching.

### Microglial phagocytosis quantification

Once microglia were isolated, their branches were examined for phagocytic cups. Phagocytic cups were defined as clear, circular morphologies appearing at the distal ends of branches. For each phagocytic event, the start and end frames were recorded, and the maximum cup diameter was measured using the *Measure Points* tool in *Imaris*. Phagocytic events were counted when a cup was unambiguously formed, regardless of outcome. This included cups that emerged at branch tips and were subsequently drawn into the soma, as well as cups that persisted for at least one frame but were not fully retracted into the soma. A subset of events were additionally classified as ‘complete’. Completed phagocytic events included events where a clear, dense, Hoechst+ nuclear fragment could be visualized within the cup, which visibly moved with the cup as the branch was retracted into the soma. For every observed phagocytic event: (1) duration (min), defined as the time from initial cup formation to event resolution; (2) maximum cup diameter (µm), defined as the largest inner diameter of the phagocytic cup; (3) phagocytic rate (count/hr), including all phagocytic events over the full recording; and (4) complete event rate (count/hr), including only the subset of events with Hoechst+ nuclear material, were recorded.

### Microglial tracking quantification

Microglial movement was quantified using the *Spots* module in *Imaris*. *Spots* were assigned to the GFP channel and placed at the center of each soma in every frame. Tracks were generated by connecting *Spots* across time using the *Track Editor*. *Spots* placement was verified in all three dimensions (X×Y×Z) to ensure accurate soma localization. One track was generated per microglia of interest. Using these tracks: (1) speed (µm·min ¹) and (2) total track length (µm) were assessed.

### Microglial morphology quantification

Microglial somas were extracted using the *Surfaces* module in *Imaris*. A ROI was created around the microglia of interest, and the *Auto Threshold* function was applied. Threshold values were verified to ensure complete capture of soma boundaries, without overfitting. *Surface* smoothing was set to 0.7 µm, and the *Seed Point* diameter was fixed at 5 µm. *Seed Points* were positioned to include one per soma and one per branch, facilitating separation of the soma from neighboring processes. If any branches remained attached to the soma, they were manually separated using the *Cut* tool. Microglial cell somas were visually distinguishable by high fluorescence intensity, underlying Hoechst staining, and often circular morphology. With microglial soma defined, the *Filaments* module was used to reconstruct processes. The smoothed soma surface was imported into the *Filament Creation Wizard* with *Auto Center* enabled. The process diameter was set to the default value of 2.10 µm to ensure consistency across samples.

### *Filaments* were created using the *Imaris AutoPath* function

The initial time-point (Frame 1) and subsequent frames (Frames 7, 14, 21, 28, 35, and 42) were selected to provide a representative sampling across the duration of the recording. This approach allowed for assessment of microglial morphology at multiple points throughout the imaging period. Using this morphological reconstruction: (1) filament length (total) (µm), the sum of all branch lengths per cell; (2) branch points, the total number of branch nodes; (3) terminal points, the total number of branch endpoints; (4) Sholl intersections, counted with a radial step size of 5 µm from the soma center; and (5) soma volume (µm³) were measured.

### Morphological clustering analysis of microglia

To compare the original 2D static morphological clustering analysis with microglial morphological attributes observed during live-cell imaging, a subset of frames from the live-cell time series were selected (e.g., Frames 7 and 35, representing early and late time-points). For each embryo, maximum-intensity projections (MIPs) were generated from selected frames using ZEN 3.8 software. These MIPs were then processed using the previously adapted protocol (70,83). Global thresholding was set to 20/255. Microglia identified as viable during live-cell imaging were analyzed. All other methodological quantifications were kept consistent with the previously published analyses (70,83).

### Quantification and statistical analysis

Statistics were represented as bar graphs showing means ± SEM or a line graph showing means ± SEM for frame-by-frame microglial movement tracking data. Live-cell imaging quantifications from the full recording were compared using two-way ANOVA with Tukey-adjusted post hoc comparisons. Time-resolved data were analyzed using a mixed-effects model with Sidak adjusted post hoc comparisons and microglia ID as a random effect. All analysis were implemented in GraphPad Prism 10.6.1.

## Declarations

### Ethics approval and consent to participate

Animal work was carried out in accordance with the guidelines and regulations of the Canadian Council on Animal Care and received prior approval from UBC’s Animal Care Committee (protocols A21-0170 and A21-0171). All experimental work was conducted in accordance with an approved Biosafety Protocol (B21-0133).

### Consent for publication

Not applicable.

### Availability of data and materials

Original images and/or videos used for analyses across the current study are available upon request to the lead contact. The datasets generated and/or analyzed during the current study are available in the GitHub repository: https://github.com/RosinLabUBC/Live-Cell-Microglia-Cold-Stress-Project-AL.git

### Authors’ contributions

JMR conceived and designed the study, assisted with fetal brain dissections, tissue sectioning, live-cell imaging, and advised on all data-analysis approaches. AL conducted mouse handling, assisted with live-cell imaging, designed the *Imaris* quantification techniques, implemented all data analysis, and conducted all statistical analysis. MR assisted with mouse handling, live-cell imaging, and microscopy. AL and JMR drafted the original manuscript, with edits incorporated from AL, MR, and JMR. All authors read and approved the final manuscript.

## Acknowledgements

The authors thank the University of British Columbia (UBC) Centre for Disease Modelling for animal care, Biljana Stojkova from the Applied Statistics and Data Science group for statistical guidance, and Erich Giedzinki from *Imaris* as well as Guang Gao from the LSI Imaging core for technical *Imaris* guidance. We also thank Mark Cembrowski and the Cembrowski lab for the use of their vibratome for our live-cell experiments. Figure 2A and Supplementary Figure 1A were created with BioRender.com. Portions of this manuscript appear in AL’s Master of Science thesis at UBC.

## Abbreviations

3D: three-dimensional
ADHD: attention-deficit/hyperactivity disorder
ANOVA: analysis of variance
ASD: autism spectrum disorder
ATP: adenosine triphosphate
AVP: arginine vasopressin
DNA: deoxyribonucleic acid
E: embryonic day
EDTA: ethylenediaminetetraacetic acid
eGFP: enhanced green fluorescent protein
MIA: maternal immune activation
NDDs: neurodevelopmental disorders
NSPCs: neural stem and progenitor cells
OXT: oxytocin
PC: principal component
PCA: principal component analysis
PCR: polymerase chain reaction
PVN: paraventricular nucleus
ROS: reactive oxygen species
SDS: sodium dodecyl sulfate
SEM: standard error of the mean
TAE: Tris-acetate-EDTA
WT: wild-type

