## Supplementary Figure 1-2 for "Live-cell imaging of fetal microglia reveals time-dependent sex differences in their responsiveness to prenatal maternal stress"

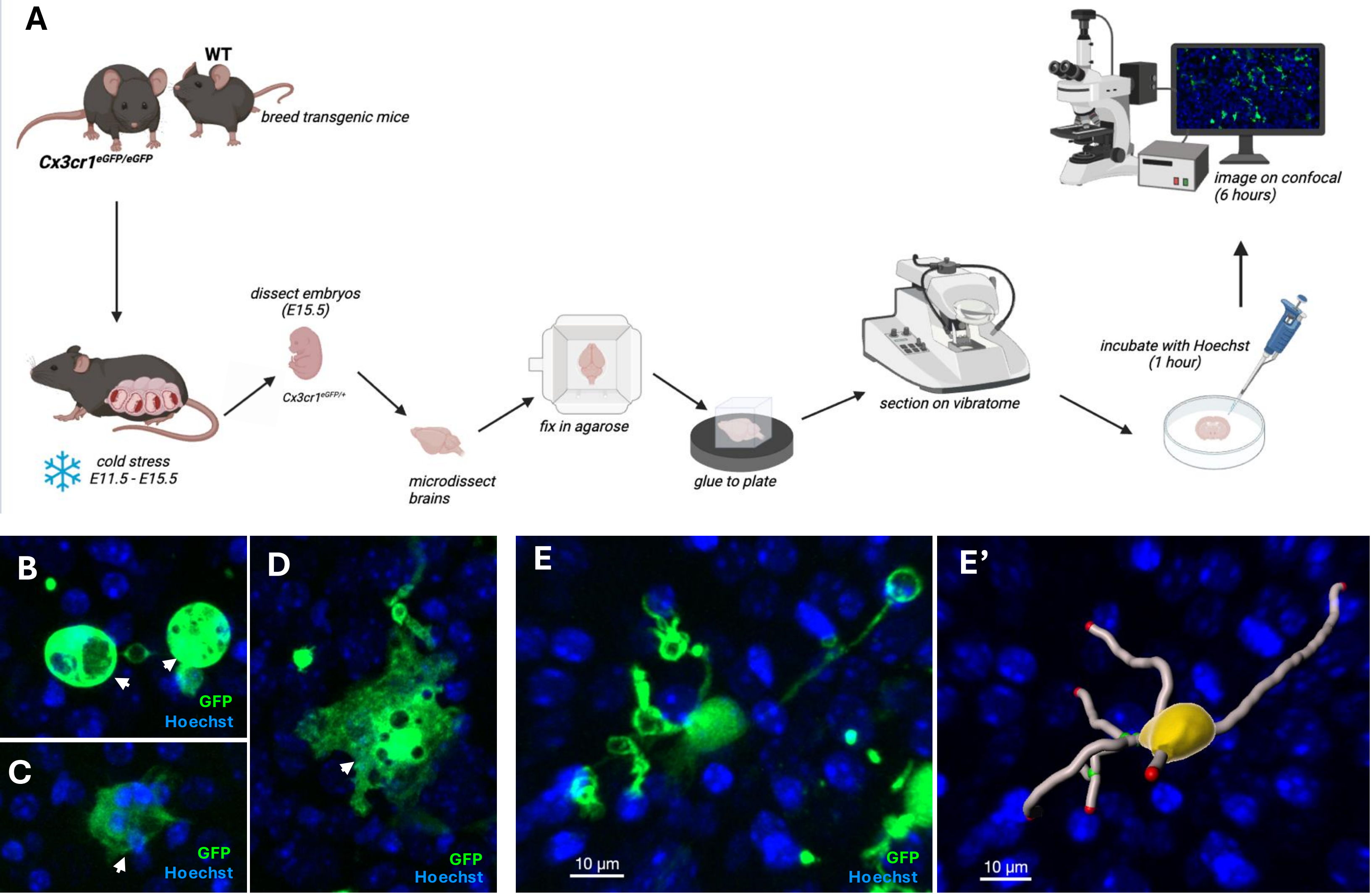

A
WT
breed transgenic mice
Cx3cr1eGFP/eGFP
cold stress
E11.5 - E15.5
dissect embryos
(E15.5)
Cx3cr1eGFP/+
microdissect
brains
fix in agarose
glue to plate
section on vibratome
incubate with Hoechst
(1 hour)
image on confocal
(6 hours)
B
GFP
Hoechst
C
GFP
Hoechst
D
GFP
Hoechst
E
10 μm
GFP
Hoechst
E'
10 μm

**Supplementary Figure 1. Live-cell imaging methodology for assessing microglial activity in the E15.5 PVN.** (A) Schematic illustrating live-cell imaging methodology used to assess microglial morphology, motility, and phagocytic activity in the E15.5 PVN. (B–D) Representative images of a microglial cell excluded from analysis due to death (B, C) and adhesion to the tissue cut surface (D). (E, E') Representative image of a microglial cell (E) and its corresponding *Imaris* 3D reconstruction (E') in the E15.5 PVN.

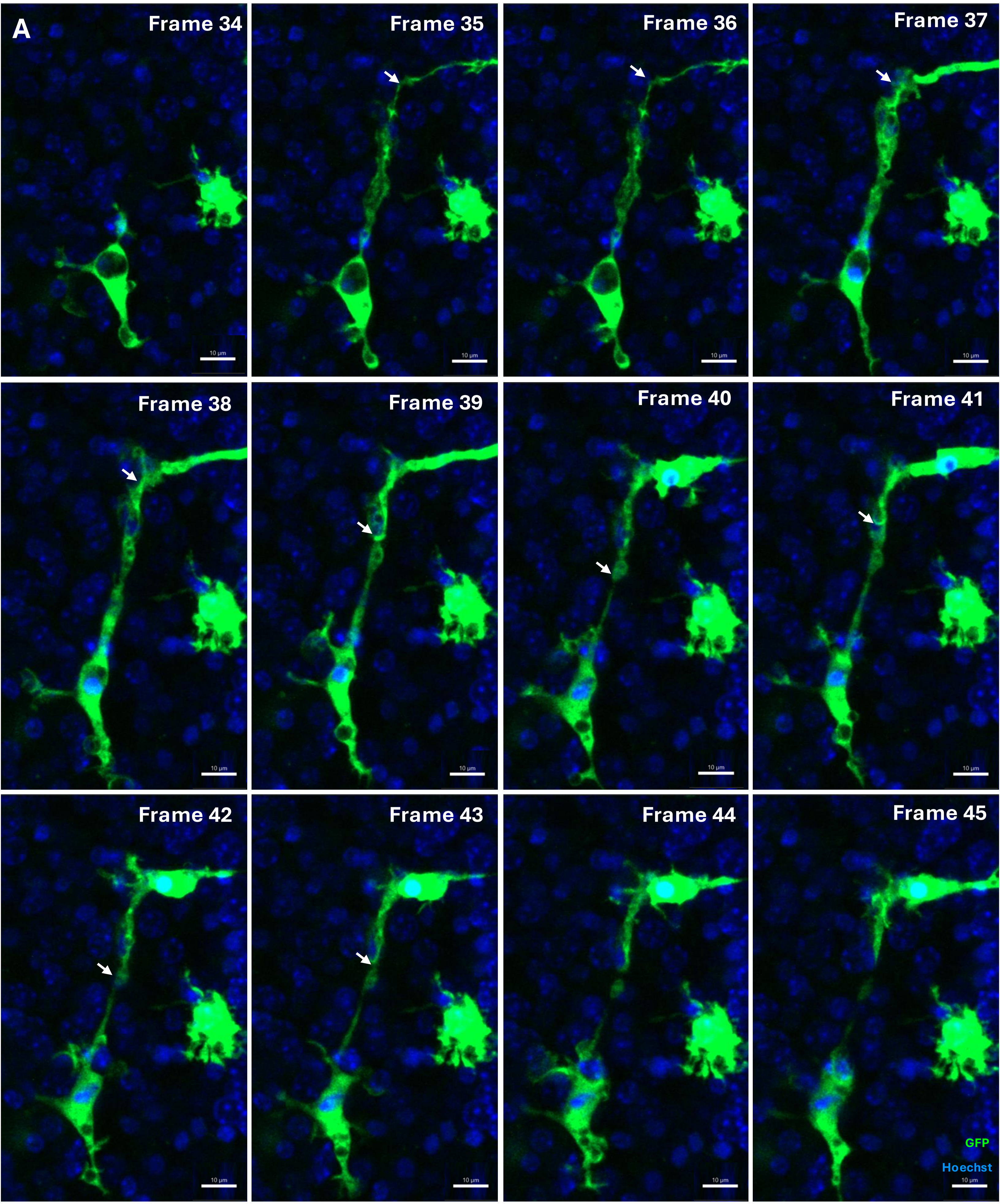
A
Frame 34
Frame 35
Frame 36
Frame 37
Frame 38
Frame 39
Frame 40
Frame 41
Frame 42
Frame 43
Frame 44
Frame 45
GFP
Hoechst

**Supplementary Figure 2. Microglia–microglia interactions during live-cell imaging in the E15.5 PVN.** (A) Sequential frames illustrating two microglial cells interacting, captured from a cold stress female embryo. White arrows indicate sites of contact.
