## Supplementary Tables 1-23 for "Live-cell imaging of fetal microglia reveals time-dependent sex differences in their responsiveness to prenatal maternal stress"

**Supplementary Table 1. Two-way ANOVA results with Tukey-adjusted post hoc for microglial soma volume in the E15.5 PVN**

| **Microglial Soma Volume** | | |
|---|---|---|
| **Tukey's multiple comparisons test** | **Adjusted p value** | **Summary** |
| Male: Control vs. Male: Cold Stress | 0.3341 | ns |
| Male: Control vs. Female: Control | 0.7016 | ns |
| Male: Control vs. Female: Cold Stress | 0.6958 | ns |
| Male: Cold Stress vs. Female: Control | 0.9088 | ns |
| Male: Cold Stress vs. Female: Cold Stress | 0.9695 | ns |
| Female: Control vs. Female: Cold Stress | 0.9990 | ns |
| **Two-way ANOVA** | **p value** | **Summary** |
| Treatment | 0.2096 | ns |
| Sex | 0.7039 | ns |
| Interaction | 0.2938 | ns |

**Supplementary Table 2. Mixed-effects ANOVA with Sidak-adjusted post hoc for microglial soma volume across time in the E15.5 male PVN**

| Microglial Soma Volume by Time in Males | | |
|---|---|---|
| **Sidak-adjusted post hoc comparisons** | **Adjusted p value** | **Summary** |
| Frame 1 (Control vs. Cold Stress) | 0.8336 | ns |
| Frame 7 (Control vs. Cold Stress) | 0.9738 | ns |
| Frame 14 (Control vs. Cold Stress) | 0.6931 | ns |
| Frame 21 (Control vs. Cold Stress) | 0.7698 | ns |
| Frame 28 (Control vs. Cold Stress) | 0.7862 | ns |
| Frame 35 (Control vs. Cold Stress) | 0.6308 | ns |
| Frame 42 (Control vs. Cold Stress) | 0.8213 | ns |
| **Mixed Effects ANOVA** | **p value** | **Summary** |
| Time | 0.0006 | *** |
| Treatment | 0.1444 | ns |
| Interaction | 0.9946 | ns |

**Supplementary Table 3. Mixed-effects ANOVA with Sidak-adjusted post hoc for microglial soma volume across time in the E15.5 female PVN**

| **Microglial Soma Volume by Time in Females** | | |
|---|---|---|
| **Sidak-adjusted post hoc comparisons** | **Adjusted p value** | **Summary** |
| Frame 1 (Control vs. Cold Stress) | 0.9486 | ns |
| Frame 7 (Control vs. Cold Stress) | 0.9158 | ns |
| Frame 14 (Control vs. Cold Stress) | 0.9997 | ns |
| Frame 21 (Control vs. Cold Stress) | 0.8934 | ns |
| Frame 28 (Control vs. Cold Stress) | 0.9958 | ns |
| Frame 35 (Control vs. Cold Stress) | 0.9992 | ns |
| Frame 42 (Control vs. Cold Stress) | 0.9998 | ns |
| **Mixed Effects ANOVA** | **p value** | **Summary** |
| Time | 0.0010 | *** |
| Treatment | 0.8134 | ns |
| Interaction | 0.1911 | ns |

**Supplementary Table 4. Two-way ANOVA results with Tukey-adjusted post hoc for microglial branch points in the E15.5 PVN**

| Microglial Branch Points | | |
|---|---|---|
| **Tukey's multiple comparisons test** | **Adjusted p value** | **Summary** |
| Male: Control vs. Male: Cold Stress | 0.4956 | ns |
| Male: Control vs. Female: Control | 0.9896 | ns |
| Male: Control vs. Female: Cold Stress | 0.1674 | ns |
| Male: Cold Stress vs. Female: Control | 0.6558 | ns |
| Male: Cold Stress vs. Female: Cold Stress | 0.8569 | ns |
| Female: Control vs. Female: Cold Stress | 0.2500 | ns |
| **Two-way ANOVA** | **p value** | **Summary** |
| Treatment | 0.0219 | * |
| Sex | 0.4252 | ns |
| Interaction | 0.6983 | ns |

**Supplementary Table 5. Mixed-effects ANOVA with Sidak-adjusted post hoc for microglial branch points across time in the E15.5 male PVN**

| Microglial Branch Points by Time in Males | | |
|---|---|---|
| **Sidak-adjusted post hoc comparisons** | **Adjusted p value** | **Summary** |
| Frame 1 (Control vs. Cold Stress) | 0.2697 | ns |
| Frame 7 (Control vs. Cold Stress) | 0.0575 | ns |
| Frame 14 (Control vs. Cold Stress) | 0.8389 | ns |
| Frame 21 (Control vs. Cold Stress) | 0.2578 | ns |
| Frame 28 (Control vs. Cold Stress) | >0.9999 | ns |
| Frame 35 (Control vs. Cold Stress) | 0.9950 | ns |
| Frame 42 (Control vs. Cold Stress) | >0.9999 | ns |
| **Mixed Effects ANOVA** | **p value** | **Summary** |
| Time | 0.0008 | *** |
| Treatment | 0.1139 | ns |
| Interaction | 0.0249 | * |

**Supplementary Table 6. Mixed-effects ANOVA with Sidak-adjusted post hoc for microglial branch points across time in the E15.5 female PVN**

| Microglial Branch Points by Time in Females | | |
|---|---|---|
| **Sidak-adjusted post hoc comparisons** | **Adjusted p value** | **Summary** |
| Frame 1 (Control vs. Cold Stress) | 0.2738 | ns |
| Frame 7 (Control vs. Cold Stress) | 0.1229 | ns |
| Frame 14 (Control vs. Cold Stress) | 0.8651 | ns |
| Frame 21 (Control vs. Cold Stress) | >0.9999 | ns |
| Frame 28 (Control vs. Cold Stress) | 0.7669 | ns |
| Frame 35 (Control vs. Cold Stress) | 0.0960 | ns |
| Frame 42 (Control vs. Cold Stress) | 0.9950 | ns |
| **Mixed Effects ANOVA** | **p value** | **Summary** |
| Time | 0.0001 | *** |
| Treatment | 0.0271 | * |
| Interaction | 0.1740 | ns |

**Supplementary Table 7. Two-way ANOVA results with Tukey-adjusted post hoc for microglial terminal points in the E15.5 PVN**

| Microglial Terminal Points | | |
|---|---|---|
| **Tukey adjusted post hoc comparisons** | **Adjusted p value** | **Summary** |
| Male: Control vs. Male: Cold Stress | 0.5472 | ns |
| Male: Control vs. Female: Control | 0.6497 | ns |
| Male: Control vs. Female: Cold Stress | 0.3190 | ns |
| Male: Cold Stress vs. Female: Control | 0.0562 | ns |
| Male: Cold Stress vs. Female: Cold Stress | 0.9563 | ns |
| Female: Control vs. Female: Cold Stress | 0.0268 | * |
| **Two-way ANOVA** | **p value** | **Summary** |
| Treatment | 0.0035 | ** |
| Sex | 0.6965 | ns |
| Interaction | 0.2499 | ns |

**Supplementary Table 8. Mixed-effects ANOVA with Sidak-adjusted post hoc for microglial terminal points across time in the E15.5 male PVN**

| Microglial Terminal Points by Time in Males | | |
|---|---|---|
| **Sidak-adjusted post hoc comparisons** | **Adjusted p value** | **Summary** |
| Frame 1 (Control vs. Cold Stress) | 0.0257 | * |
| Frame 7 (Control vs. Cold Stress) | 0.0747 | ns |
| Frame 14 (Control vs. Cold Stress) | >0.9999 | ns |
| Frame 21 (Control vs. Cold Stress) | 0.9201 | ns |
| Frame 28 (Control vs. Cold Stress) | 0.8548 | ns |
| Frame 35 (Control vs. Cold Stress) | >0.9999 | ns |
| Frame 42 (Control vs. Cold Stress) | >0.9999 | ns |
| **Mixed Effects ANOVA** | **p value** | **Summary** |
| Time | 0.3271 | ns |
| Treatment | 0.1302 | ns |
| Interaction | 0.0107 | * |

**Supplementary Table 9. Mixed-effects ANOVA with Sidak-adjusted post hoc for microglial terminal points across time in the E15.5 female PVN**

| Microglial Terminal Points by Time in Females | | |
|---|---|---|
| **Sidak-adjusted post hoc comparisons** | **Adjusted p value** | **Summary** |
| Frame 1 (Control vs. Cold Stress) | 0.9946 | ns |
| Frame 7 (Control vs. Cold Stress) | 0.3630 | ns |
| Frame 14 (Control vs. Cold Stress) | 0.1661 | ns |
| Frame 21 (Control vs. Cold Stress) | 0.4856 | ns |
| Frame 28 (Control vs. Cold Stress) | 0.0484 | * |
| Frame 35 (Control vs. Cold Stress) | 0.0179 | * |
| Frame 42 (Control vs. Cold Stress) | 0.1238 | ns |
| **Mixed Effects ANOVA** | **p value** | **Summary** |
| Time | 0.5502 | ns |
| Treatment | 0.0040 | ** |
| Interaction | 0.7445 | ns |

**Supplementary Table 10. Two-way ANOVA with Tukey-adjusted post hoc for microglial Sholl intersections in the E15.5 PVN**

| **Average Microglial Sholl Intersections** | | |
|---|---|---|
| **Tukey-adjusted post hoc comparisons** | **Adjusted p value** | **Summary** |
| Male: Control vs. Male: Cold Stress | 0.5937 | ns |
| Male: Control vs. Female: Control | 0.9932 | ns |
| Male: Control vs. Female: Cold Stress | 0.0367 | * |
| Male: Cold Stress vs. Female: Control | 0.3954 | ns |
| Male: Cold Stress vs. Female: Cold Stress | 0.3881 | ns |
| Female: Control vs. Female: Cold Stress | 0.0142 | * |
| **Two-way ANOVA** | **p value** | **Summary** |
| Treatment | 0.0025 | ** |
| Sex | 0.3140 | ns |
| Interaction | 0.1739 | ns |

**Supplementary Table 11. Mixed-effects ANOVA with Sidak-adjusted post hoc for microglial Sholl intersections across time in the E15.5 male PVN**

| Microglial Sholl Intersections by Time in Males | | |
|---|---|---|
| **Sidak-adjusted post hoc comparisons** | **Adjusted p value** | **Summary** |
| Frame 1 (Control vs. Cold Stress) | 0.2119 | ns |
| Frame 7 (Control vs. Cold Stress) | 0.0276 | * |
| Frame 14 (Control vs. Cold Stress) | 0.9747 | ns |
| Frame 21 (Control vs. Cold Stress) | 0.7839 | ns |
| Frame 28 (Control vs. Cold Stress) | 0.9941 | ns |
| Frame 35 (Control vs. Cold Stress) | >0.9999 | ns |
| Frame 42 (Control vs. Cold Stress) | 0.9998 | ns |
| **Mixed Effects ANOVA** | **p value** | **Summary** |
| Time | <0.0001 | **** |
| Treatment | 0.1480 | ns |
| Interaction | 0.0193 | * |

**Supplementary Table 12. Mixed-effects ANOVA with Sidak-adjusted post hoc for microglial Sholl intersections across time in the E15.5 female PVN**

| **Microglial Sholl Intersections by Time in Females** | | |
|---|---|---|
| **Sidak-adjusted post hoc comparisons** | **Adjusted p value** | **Summary** |
| Frame 1 (Control vs. Cold Stress) | 0.0537 | ns |
| Frame 7 (Control vs. Cold Stress) | 0.4186 | ns |
| Frame 14 (Control vs. Cold Stress) | 0.0795 | ns |
| Frame 21 (Control vs. Cold Stress) | 0.6460 | ns |
| Frame 28 (Control vs. Cold Stress) | 0.0156 | * |
| Frame 35 (Control vs. Cold Stress) | 0.0461 | * |
| Frame 42 (Control vs. Cold Stress) | 0.0588 | ns |
| **Mixed Effects ANOVA** | **p value** | **Summary** |
| Time | 0.0003 | *** |
| Treatment | 0.0040 | ** |
| Interaction | 0.4227 | ns |

**Supplementary Table 13. Two-way ANOVA results with Tukey-adjusted post hoc for microglial branch length in the E15.5 PVN**

| **Microglial Branch Length** | | |
|---|---|---|
| **Tukey-adjusted post hoc comparisons** | **Adjusted p value** | **Summary** |
| Male: Control vs. Male: Cold Stress | 0.5134 | ns |
| Male: Control vs. Female: Control | 0.9946 | ns |
| Male: Control vs. Female: Cold Stress | 0.0236 | * |
| Male: Cold Stress vs. Female: Control | 0.6352 | ns |
| Male: Cold Stress vs. Female: Cold Stress | 0.3626 | ns |
| Female: Control vs. Female: Cold Stress | 0.0336 | * |
| **Two-way ANOVA** | **p value** | **Summary** |
| Treatment | 0.0038 | ** |
| Sex | 0.1676 | ns |
| Interaction | 0.2917 | ns |

**Supplementary Table 14. Mixed-effects ANOVA with Sidak-adjusted post hoc for microglial branch length across time in the E15.5 male PVN**

| **Microglial Branch Length by Time in Males** | | |
|---|---|---|
| **Sidak-adjusted post hoc comparisons** | **Adjusted p value** | **Summary** |
| Frame 1 (Control vs. Cold Stress) | 0.1136 | ns |
| Frame 7 (Control vs. Cold Stress) | 0.0308 | * |
| Frame 14 (Control vs. Cold Stress) | 0.6784 | ns |
| Frame 21 (Control vs. Cold Stress) | 0.6566 | ns |
| Frame 28 (Control vs. Cold Stress) | 0.9848 | ns |
| Frame 35 (Control vs. Cold Stress) | >0.9999 | ns |
| Frame 42 (Control vs. Cold Stress) | >0.9999 | ns |
| **Mixed Effects ANOVA** | **p value** | **Summary** |
| Time | <0.0001 | **** |
| Treatment | 0.1020 | ns |
| Interaction | 0.0182 | * |

**Supplementary Table 15. Mixed-effects ANOVA with Sidak-adjusted post hoc for microglial branch length across time in the E15.5 female PVN**

| **Microglial Branch Length by Time in Females** | | |
|---|---|---|
| **Sidak-adjusted post hoc comparisons** | **Adjusted p value** | **Summary** |
| Frame 1 (Control vs. Cold Stress) | 0.2336 | ns |
| Frame 7 (Control vs. Cold Stress) | 0.3813 | ns |
| Frame 14 (Control vs. Cold Stress) | 0.1019 | ns |
| Frame 21 (Control vs. Cold Stress) | 0.6674 | ns |
| Frame 28 (Control vs. Cold Stress) | 0.0664 | ns |
| Frame 35 (Control vs. Cold Stress) | 0.1200 | ns |
| Frame 42 (Control vs. Cold Stress) | 0.0824 | ns |
| **Mixed Effects ANOVA** | **p value** | **Summary** |
| Time | <0.0001 | **** |
| Treatment | 0.0099 | ** |
| Interaction | 0.7570 | ns |

**Supplementary Table 16. Two-way ANOVA results with Tukey-adjusted post hoc for total microglial distance in the E15.5 PVN**

| **Average Distance Traveled** | | |
|---|---|---|
| **Tukey-adjusted post hoc comparisons** | **Adjusted p value** | **Summary** |
| Male: Control vs. Male: Cold Stress | 0.8693 | ns |
| Male: Control vs. Female: Control | 0.1208 | ns |
| Male: Control vs. Female: Cold Stress | 0.0122 | * |
| Male: Cold Stress vs. Female: Control | 0.4712 | ns |
| Male: Cold Stress vs. Female: Cold Stress | 0.0769 | ns |
| Female: Control vs. Female: Cold Stress | 0.6374 | ns |
| **Two-way ANOVA** | **p value** | **Summary** |
| Treatment | 0.1675 | ns |
| Sex | 0.0013 | ** |
| Interaction | 0.7367 | ns |

**Supplementary Table 17. Two-way ANOVA results with Tukey-adjusted post hoc for microglial speed in the E15.5 PVN**

| **Average Microglial Speed** | | |
|---|---|---|
| **Tukey-adjusted post hoc comparisons** | **Adjusted p value** | **Summary** |
| Male: Control vs. Male: Cold Stress | 0.9984 | ns |
| Male: Control vs. Female: Control | 0.5445 | ns |
| Male: Control vs. Female: Cold Stress | 0.0024 | ** |
| Male: Cold Stress vs. Female: Control | 0.4331 | ns |
| Male: Cold Stress vs. Female: Cold Stress | 0.0013 | ** |
| Female: Control vs. Female: Cold Stress | 0.0608 | ns |
| **Two-way ANOVA** | **p value** | **Summary** |
| Treatment | 0.0867 | ns |
| Sex | 0.0003 | *** |
| Interaction | 0.0534 | ns |

**Supplementary Table 18. Mixed-effects ANOVA results with Sidak-adjusted post hoc for microglial speed in the E15.5 male PVN**

| Microglial Speed in Males | | |
|---|---|---|
| **Sidak-adjusted post hoc comparisons** | **Adjusted p value** | **Summary** |
| Frame 1 (Control vs. Cold Stress) | 0.0796 | ns |
| Frame 2 (Control vs. Cold Stress) | 0.7721 | ns |
| Frame 3 (Control vs. Cold Stress) | >0.9999 | ns |
| Frame 4 (Control vs. Cold Stress) | >0.9999 | ns |
| Frame 5 (Control vs. Cold Stress) | >0.9999 | ns |
| Frame 6 (Control vs. Cold Stress) | >0.9999 | ns |
| Frame 7 (Control vs. Cold Stress) | 0.9967 | ns |
| Frame 8 (Control vs. Cold Stress) | 0.9886 | ns |
| Frame 9 (Control vs. Cold Stress) | >0.9999 | ns |
| Frame 10 (Control vs. Cold Stress) | 0.9967 | ns |
| Frame 11 (Control vs. Cold Stress) | 0.8998 | ns |
| Frame 12 (Control vs. Cold Stress) | 0.8908 | ns |
| Frame 13 (Control vs. Cold Stress) | >0.9999 | ns |
| Frame 14 (Control vs. Cold Stress) | >0.9999 | ns |
| Frame 15 (Control vs. Cold Stress) | >0.9999 | ns |
| Frame 16 (Control vs. Cold Stress) | >0.9999 | ns |
| Frame 17 (Control vs. Cold Stress) | >0.9999 | ns |
| Frame 18 (Control vs. Cold Stress) | >0.9999 | ns |
| Frame 19 (Control vs. Cold Stress) | >0.9999 | ns |
| Frame 20 (Control vs. Cold Stress) | >0.9999 | ns |
| Frame 21 (Control vs. Cold Stress) | >0.9999 | ns |
| Frame 22 (Control vs. Cold Stress) | 0.9862 | ns |
| Frame 23 (Control vs. Cold Stress) | >0.9999 | ns |
| Frame 24 (Control vs. Cold Stress) | >0.9999 | ns |
| Frame 25 (Control vs. Cold Stress) | >0.9999 | ns |
| Frame 26 (Control vs. Cold Stress) | >0.9999 | ns |
| Frame 27 (Control vs. Cold Stress) | >0.9999 | ns |
| Frame 28 (Control vs. Cold Stress) | >0.9999 | ns |
| Frame 29 (Control vs. Cold Stress) | >0.9999 | ns |
| Frame 30 (Control vs. Cold Stress) | >0.9999 | ns |
| Frame 31 (Control vs. Cold Stress) | >0.9999 | ns |
| Frame 32 (Control vs. Cold Stress) | >0.9999 | ns |
| Frame 33 (Control vs. Cold Stress) | 0.9666 | ns |
| Frame 34 (Control vs. Cold Stress) | >0.9999 | ns |

| | | |
|---|---|---|
| Frame 35 (Control vs. Cold Stress) | >0.9999 | ns |
| Frame 36 (Control vs. Cold Stress) | >0.9999 | ns |
| Frame 37 (Control vs. Cold Stress) | >0.9999 | ns |
| Frame 38 (Control vs. Cold Stress) | 0.1302 | ns |
| Frame 39 (Control vs. Cold Stress) | 0.3211 | ns |
| Frame 40 (Control vs. Cold Stress) | >0.9999 | ns |
| Frame 41 (Control vs. Cold Stress) | >0.9999 | ns |
| Frame 42 (Control vs. Cold Stress) | >0.9999 | ns |
| Frame 43 (Control vs. Cold Stress) | >0.9999 | ns |
| Frame 44 (Control vs. Cold Stress) | >0.9999 | ns |
| Frame 45 (Control vs. Cold Stress) | >0.9999 | ns |
| **Mixed Effects ANOVA** | **p value** | **Summary** |
| Time | <0.0001 | **** |
| Treatment | 0.5070 | ns |
| Interaction | 0.0023 | ** |

**Supplementary Table 19. Mixed-effects ANOVA results with Sidak-adjusted post hoc for microglial speed in the E15.5 female PVN**

| Microglial Speed in Females | | |
|---|---|---|
| **Sidak-adjusted post hoc comparisons** | **Adjusted p value** | **Summary** |
| Frame 1 (Control vs. Cold Stress) | <0.0001 | **** |
| Frame 2 (Control vs. Cold Stress) | 0.0009 | *** |
| Frame 3 (Control vs. Cold Stress) | 0.8434 | ns |
| Frame 4 (Control vs. Cold Stress) | 0.9858 | ns |
| Frame 5 (Control vs. Cold Stress) | 0.9994 | ns |
| Frame 6 (Control vs. Cold Stress) | >0.9999 | ns |
| Frame 7 (Control vs. Cold Stress) | 0.9891 | ns |
| Frame 8 (Control vs. Cold Stress) | 0.2388 | ns |
| Frame 9 (Control vs. Cold Stress) | >0.9999 | ns |
| Frame 10 (Control vs. Cold Stress) | >0.9999 | ns |
| Frame 11 (Control vs. Cold Stress) | 0.7292 | ns |
| Frame 12 (Control vs. Cold Stress) | >0.9999 | ns |
| Frame 13 (Control vs. Cold Stress) | >0.9999 | ns |
| Frame 14 (Control vs. Cold Stress) | 0.9956 | ns |
| Frame 15 (Control vs. Cold Stress) | >0.9999 | ns |
| Frame 16 (Control vs. Cold Stress) | 0.9666 | ns |
| Frame 17 (Control vs. Cold Stress) | >0.9999 | ns |
| Frame 18 (Control vs. Cold Stress) | >0.9999 | ns |
| Frame 19 (Control vs. Cold Stress) | >0.9999 | ns |
| Frame 20 (Control vs. Cold Stress) | >0.9999 | ns |
| Frame 21 (Control vs. Cold Stress) | 0.9972 | ns |
| Frame 22 (Control vs. Cold Stress) | >0.9999 | ns |
| Frame 23 (Control vs. Cold Stress) | >0.9999 | ns |
| Frame 24 (Control vs. Cold Stress) | 0.9997 | ns |
| Frame 25 (Control vs. Cold Stress) | 0.9932 | ns |
| Frame 26 (Control vs. Cold Stress) | >0.9999 | ns |
| Frame 27 (Control vs. Cold Stress) | >0.9999 | ns |
| Frame 28 (Control vs. Cold Stress) | >0.9999 | ns |
| Frame 29 (Control vs. Cold Stress) | >0.9999 | ns |
| Frame 30 (Control vs. Cold Stress) | >0.9999 | ns |
| Frame 31 (Control vs. Cold Stress) | >0.9999 | ns |
| Frame 32 (Control vs. Cold Stress) | 0.9841 | ns |
| Frame 33 (Control vs. Cold Stress) | >0.9999 | ns |
| Frame 34 (Control vs. Cold Stress) | >0.9999 | ns |

| | | |
|---|---|---|
| Frame 35 (Control vs. Cold Stress) | >0.9999 | ns |
| Frame 36 (Control vs. Cold Stress) | 0.9356 | ns |
| Frame 37 (Control vs. Cold Stress) | 0.6940 | ns |
| Frame 38 (Control vs. Cold Stress) | >0.9999 | ns |
| Frame 39 (Control vs. Cold Stress) | >0.9999 | ns |
| Frame 40 (Control vs. Cold Stress) | >0.9999 | ns |
| Frame 41 (Control vs. Cold Stress) | >0.9999 | ns |
| Frame 42 (Control vs. Cold Stress) | >0.9999 | ns |
| Frame 43 (Control vs. Cold Stress) | >0.9999 | ns |
| Frame 44 (Control vs. Cold Stress) | >0.9999 | ns |
| Frame 45 (Control vs. Cold Stress) | >0.9999 | ns |
| **Mixed Effects ANOVA** | **p value** | **Summary** |
| Time | <0.0001 | **** |
| Treatment | 0.0080 | ** |
| Interaction | 0.0023 | ** |

**Supplementary Table 20. Two-way ANOVA results with Tukey-adjusted post hoc for phagocytic cup diameter in the E15.5 PVN**

| Microglial Phagocytic Cup Diameter | | |
|---|---|---|
| **Tukey-adjusted post hoc comparisons** | **Adjusted p value** | **Summary** |
| Male: Control vs. Male: Cold Stress | 0.1536 | ns |
| Male: Control vs. Female: Control | >0.9999 | ns |
| Male: Control vs. Female: Cold Stress | 0.9413 | ns |
| Male: Cold Stress vs. Female: Control | 0.1266 | ns |
| Male: Cold Stress vs. Female: Cold Stress | 0.5258 | ns |
| Female: Control vs. Female: Cold Stress | 0.9257 | ns |
| **Two-way ANOVA** | **p value** | **Summary** |
| Treatment | 0.0609 | ns |
| Sex | 0.2997 | ns |
| Interaction | 0.3280 | ns |

**Supplementary Table 21. Two-way ANOVA results with Tukey-adjusted post hoc for phagocytic event duration in the E15.5 PVN**

| **Microglial Phagocytic Event Duration** | | |
|---|---|---|
| **Tukey-adjusted post hoc comparisons** | **Adjusted p value** | **Summary** |
| Male: Control vs. Male: Cold Stress | 0.5035 | ns |
| Male: Control vs. Female: Control | 0.8499 | ns |
| Male: Control vs. Female: Cold Stress | 0.6793 | ns |
| Male: Cold Stress vs. Female: Control | 0.9278 | ns |
| Male: Cold Stress vs. Female: Cold Stress | 0.9988 | ns |
| Female: Control vs. Female: Cold Stress | 0.9781 | ns |
| **Two-way ANOVA** | **p value** | **Summary** |
| Treatment | 0.2164 | ns |
| Sex | 0.6686 | ns |
| Interaction | 0.5143 | ns |

**Supplementary Table 22. Two-way ANOVA results with Tukey-adjusted post hoc for microglial phagocytic events in the E15.5 PVN**

| Microglial Phagocytic Events (per hour) | | |
|---|---|---|
| **Tukey's multiple comparisons test** | | |
| **Tukey-adjusted post hoc comparisons** | **Adjusted p value** | **Summary** |
| Male: Control vs. Male: Cold Stress | 0.6255 | ns |
| Male: Control vs. Female: Control | 0.0195 | * |
| Male: Control vs. Female: Cold Stress | 0.5671 | ns |
| Male: Cold Stress vs. Female: Control | 0.3168 | ns |
| Male: Cold Stress vs. Female: Cold Stress | 0.0808 | ns |
| Female: Control vs. Female: Cold Stress | 0.0005 | *** |
| **Two-way ANOVA** | **p value** | **Summary** |
| Treatment | 0.0369 | * |
| Sex | 0.8748 | ns |
| Interaction | 0.0003 | *** |

**Supplementary Table 23. Two-way ANOVA results with Tukey-adjusted post hoc for completed phagocytic events in the E15.5 PVN**

| Microglial Complete Phagocytic Events (per hour) | | |
|---|---|---|
| **Tukey-adjusted post hoc comparisons** | **Adjusted p value** | **Summary** |
| Male: Control vs. Male: Cold Stress | 0.0034 | ** |
| Male: Control vs. Female: Control | 0.0016 | ** |
| Male: Control vs. Female: Cold Stress | 0.1518 | ns |
| Male: Cold Stress vs. Female: Control | 0.9996 | ns |
| Male: Cold Stress vs. Female: Cold Stress | 0.6982 | ns |
| Female: Control vs. Female: Cold Stress | 0.6192 | ns |
| **Two-way ANOVA** | **p value** | **Summary** |
| Treatment | 0.1294 | ns |
| Sex | 0.0978 | ns |
| Interaction | 0.0013 | ** |
